# Downstream of gasdermin D cleavage, a Ragulator-Rag-mTORC1 pathway promotes pore formation and pyroptosis

**DOI:** 10.1101/2020.11.02.362517

**Authors:** Charles L. Evavold, Iva Hafner-Bratkovič, Jonathan C. Kagan

**Author notes:** These authors contributed equally (Charles L. Evavold, Iva Hafner-Bratkovič).

## Abstract

The process of pyroptosis is mediated by inflammasomes and a downstream effector known as gasdermin D (GSDMD). Upon cleavage by inflammasome-associated caspases, the N-terminal domain of GSDMD forms membrane pores that promote cytolysis. Numerous proteins are recognized to promote GSDMD cleavage, but none are known to be required for pore formation after GSDMD cleavage. Herein, we report a forward genetic screen that was designed to identify regulators of pyroptosis that act downstream of GSDMD cleavage. This screen identified several components of the Ragulator-Rag complex, which is known for its metabolic control of mTOR. Mechanistic studies revealed that Ragulator-Rag is not necessary for GSDMD localization to the plasma membrane, but is necessary for pore formation and mitochondrial inactivation. Downstream of Ragulator-Rag is mTORC1, which we found to promote pyroptosis in response to diverse natural stimuli, including infection. GSDMD therefore requires a Ragulator-Rag-mTORC1 pathway in order to form pores and execute pyroptosis.

## Introduction

The link between cell death and inflammation has long-been recognized, with certain death processes inducing immuno-suppressive responses (*e.g.* apoptosis) and others inducing inflammatory responses (*e.g.* pyroptosis). Central to the inflammatory process of pyroptosis is the protein gasdermin D (GSDMD) (Kayagaki et al., 2015; Shi et al., 2015), which forms large pores in the plasma membrane of cells that can result in lysis and the release of intracellular inflammatory mediators (Kovacs and Miao, 2017; Lieberman et al., 2019). Among these inflammatory mediators are interleukin (IL)-1 family cytokines (Dinarello, 2018), which are cytosolic proteins that can be released into the extracellular space after GSDMD pore formation from living (Evavold et al., 2018; Heilig et al., 2018) or pyroptotic cells (He et al., 2015; Kayagaki et al., 2015; Shi et al., 2015). In contrast to their pyroptotic counterparts, apoptotic cells typically maintain plasma membrane integrity and therefore do not release intracellular inflammatory mediators. Distinctions in plasma membrane integrity therefore (in part) help explain inflammatory or non-inflammatory consequences of the different processes of cell death.

Two well-established mechanisms explain how pyroptosis-inducing signal transduction pathways stimulate GSDMD pore formation and inflammation. The first mechanism involves the actions of inflammasomes, which are supramolecular organizing centers that function as the subcellular sites of caspase-1 activation (Chan and Schroder, 2020; Samir and Kanneganti, 2019). Caspase-1 is a dormant enzyme in resting cells. Upon infection or select disruptions of cellular homeostasis, inflammasomes are assembled in the cytosol that recruit and activate caspase-1. This activated enzyme cleaves GSDMD into two fragments, the N-terminus of which oligomerizes and inserts into the plasma membrane where it forms pores of 10-20 nm in inner diameter (Aglietti et al., 2016; Ding et al., 2016; Liu et al., 2016; Ruan et al., 2018; Sborgi et al., 2016). Caspase-1 also cleaves the pro-IL-1 family members IL-1β and IL-18 into their bioactive inflammatory forms, which are then released from cells after GSDMD pores are formed. The second means by which pyroptosis is induced is by the actions of murine caspase-11 (or caspase-4 and −5 in humans). The catalytic activity of these caspases is not stimulated upon recruitment into inflammasomes. Rather, catalytic activity is stimulated upon binding to bacterial lipopolysaccharides (LPS) that reach the cytosol (Hagar et al., 2013; Kayagaki et al., 2013; Shi et al., 2014). Upon LPS binding, active caspase-11 cleaves GSDMD in a manner similar to caspase-1 (Wang et al., 2020), leading to pore formation and pyroptosis (Aglietti et al., 2016; Kayagaki et al., 2015; Liu et al., 2016; Shi et al., 2015). Other mechanisms of pyroptosis are emerging, which are mediated by distinct caspases or GSDM family members (Orning et al., 2018; Rogers et al., 2017; Wang et al., 2017).

Despite the widespread appreciation of the importance of GSDMD in pyroptosis, mechanisms regulating its activity are largely focused on upstream factors that influence its cleavage. For example, genetic deficiencies in proteins that seed inflammasome assembly (*e.g.* NLRP3) or core components of inflammasomes (*e.g.* ASC) prevent caspase-1 activation in response to numerous infectious encounters or disruptions of cellular homeostasis. Similar to deficiencies of the caspases that cleave GSDMD, NLRP3 or ASC deficiencies result in inefficient GSDMD cleavage and defects in pyroptosis. Moreover, certain stimuli and cell types may differentially impact the magnitude and duration of caspase activation, with a direct effect on the rate of GSDMD cleavage and pore formation (Boucher et al., 2018; Chen et al., 2014). Recent work indicates that TCA cycle intermediates modify a reactive cysteine within GSDMD resulting in reduced cleavage and pore formation (Humphries et al., 2020). While the above-mentioned mechanisms of regulation are diverse, they all operate upstream of GSDMD cleavage. Downstream of GSDMD cleavage, membrane repair pathways act to limit the extent of GSDMD pore formation at the plasma membrane (Ruhl et al., 2018). Whether factors exist that promote pore formation at the plasma membrane after GSDMD cleavage has occurred is less clear.

In this study, we sought to fill this gap in our knowledge by performing a genome-wide forward genetic screen to identify factors that act downstream of GSDMD cleavage, which are required for pore forming activity at the plasma membrane. We identified multiple genes that are necessary for GSDMD pore forming activity in macrophages. These genes were highly enriched in factors that encode distinct components of the Ragulator-Rag complex, a central mediator of mTOR-dependent activities. Subsequent mechanistic and functional analysis revealed that the Ragulator-Rag complex controls mTORC1 (but not mTORC2)-dependent events that operate downstream of GSDMD cleavage and membrane localization to promote pore formation and pyroptosis.

## Results

### An experimental platform that bypasses natural regulatory mechanisms to specifically stimulate GSDMD pore forming activity

Several genetic screens have been performed to identify regulators of pyroptosis (Kayagaki et al., 2019; Kayagaki et al., 2015; Napier et al., 2016; Shi et al., 2015). These screens relied on cell stimulations that activated signaling pathways upstream of GSDMD cleavage and pore formation. Consequently, the genes identified were found to function at multiple stages of the pyroptotic process. We sought to create an experimental screening system that would focus on the specific stage that GSDMD operates within pyroptotic pathways. This was accomplished by taking advantage of the fact that the expression of the N-terminal caspase-cleavage product of GSDMD is sufficient to induce pore formation in the plasma membrane (Evavold et al., 2018; Kayagaki et al., 2015; Shi et al., 2015). Thus, expression of this N-terminal GSDMD fragment (NT-GSDMD) bypasses all upstream regulatory events that naturally lead to pyroptosis.

To identify genes that are necessary for plasma membrane permeabilization induced by NT-GSDMD, we established a doxycycline (Dox)-inducible system suitable for forward genetic analysis. Using immortalized bone marrow-derived macrophages (iBMDMs) from Cas9-expressing transgenic mice, we stably expressed the Tet3G transactivator gene, which enabled the Dox-inducible expression of a transgene of interest. Dox-inducible transgenes encoding NT-GSDMD and full length GSDMD (FL-GSDMD) were introduced into these cells, both of which contained a C-terminal BFP tag (Figure 1A and 1B). We utilized a variant of the NT-GSDMD that contains an I105N mutation, as this mutant has been shown by us and others to be more readily detected within cells than their wild type (WT) counterpart (Aglietti et al., 2016; Evavold et al., 2018). Importantly, this mutant forms pores in the plasma membrane similar to WT GSDMD, albeit less efficiently.

**Figure 1:**
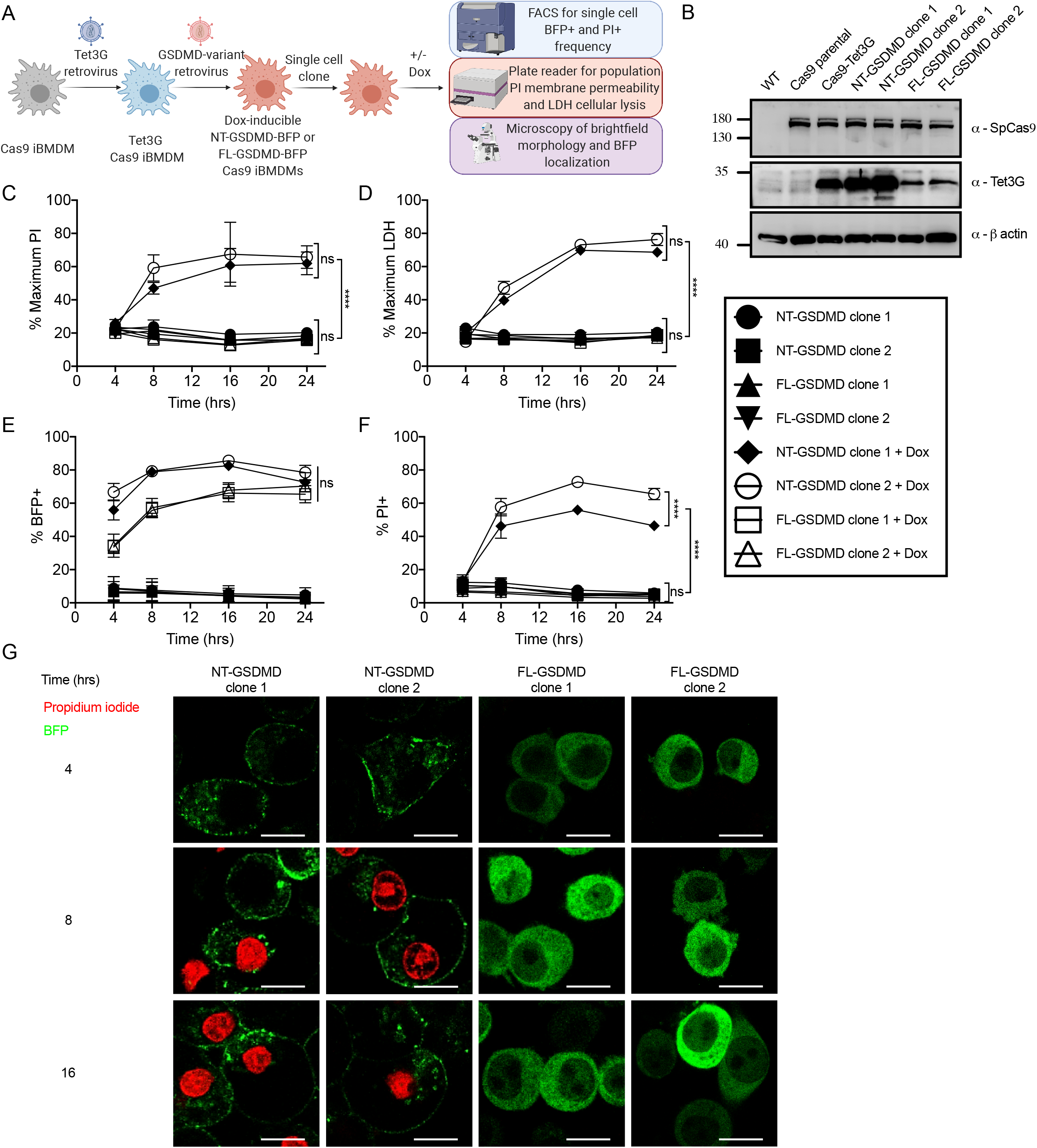
Synthetically engineered macrophages model the execution phase of pyroptosis through direct production of the pore forming N terminus of GSDMD. (A) Retroviral transduction work flow to generate stable Tet3G transactivator expressing doxycycline inducible fluorescently tagged variants of GSDMD in iBMDMs from the Cas9 knockin mouse and downstream characterization. (B) Western blot of stable Cas9 expression in parental and progeny iBMDMs and stable expression of Tet3G transactivator in progeny iBMDM clones. (C) Kinetic analysis of PI uptake by plate reader to measure bulk membrane permeability in populations of uninduced or doxycycline induced (0.5 μg/ml) clones expressing NT-GSDMD-BFP or FL-GSDMD-BFP. (D) Time course end point analysis of LDH release into cell free supernatants to measure bulk cellular lysis in populations of uninduced or doxycycline induced (0.5 μg/ml) clones expressing NT-GSDMD-BFP or FL-GSDMD-BFP. (E) Time course end point analysis of the frequency of BFP+ cells by flow cytometry of cells uninduced or doxycycline induced (0.5 μg/ml) clones expressing NT-GSDMD-BFP or FL-GSDMD-BFP. (F) Time course end point analysis of the frequency of PI+ cells by flow cytometry of cells uninduced or doxycycline induced (0.5 μg/ml) clones expressing NT-GSDMD-BFP or FL-GSDMD-BFP. (G) Time course live cell imaging of doxycycline induced clones expressing NT-GSDMD-BFP or FL-GSDMD-BFP noting localization of BFP signal and PI uptake. Scale bar indicates 10 μm. All quantification represents the mean and SEM of three independent experiments.

We characterized the ability of NT-GSDMD and FL-GSDMD to induce plasma membrane pore formation and lysis after Dox treatment. Pore formation was assessed by treatment of cells with propidium iodide (PI), a membrane impermeable dye that can pass through GSDMD pores and enter cells. PI fluoresces upon binding to intracellular nucleic acids. Cytolysis was assessed by the release of the cytosolic enzyme lactate dehydrogenase (LDH) into the extracellular space. Unlike PI, enzymatically active LDH is too large to pass through GSDMD pores (Evavold et al., 2018). LDH is only released from cells post-lysis.

We found that Dox-mediated induction of NT-GSDMD resulted in pore formation and cell lysis, as revealed by population level PI fluorescence and LDH release (Figure 1C and 1D). In contrast, Dox-induced expression of FL-GSDMD did not result in pore formation or cell lysis (Figure 1C and 1D). The BFP fusion to each GSDMD cDNA enabled us to use flow cytometry to assess BFP fluorescence as a quantitative assessment of GSDMD expression after Dox-treatment. Induction of both GSDMD transgenes led to robust BFP induction, as evidenced by an increase in frequency of BFP positive cells over time in a Dox-dependent manner (Figure 1E). Flow cytometry also demonstrated that the frequency of PI-positive cells increased over time in a Dox-dependent manner for cells expressing NT-GSDMD, while PI positivity remained low for cells expressing FL-GSDMD (Figure 1F). We used confocal microscopy to examine GSDMD localization after Dox-induction. This analysis demonstrated that NT-GSDMD is localized primarily to the plasma membrane, whereas FL-GSDMD was distributed throughout the cell (Figure 1G). Live cell microscopy imaging demonstrated that PI staining occurred in a time-dependent manner in cells expressing the NT-GSDMD, but not in cells expressing the cytosolic FL-GSDMD (Figure 1G).

Finally, we sought to determine if NT-GSDMD mediated pyroptosis induced by our Dox system was impacted by the endogenous inflammasome machinery within the cell. This possibility was minimized throughout our study by performing all experiments in the absence of the typical LPS-priming step that is important for inflammasome activities (Bauernfeind et al., 2009; Juliana et al., 2012; O’Connor et al., 2003). Additionally, we found that Dox-induced pore forming activity of NT-GSDMD remained intact in cells treated with the NLRP3-specific inhibitor MCC950 (Coll et al., 2019; Tapia-Abellan et al., 2019) (Figure 2A and 2B). Even when cells were primed with LPS, Dox-induced NT-GSDMD mediated LDH release and PI uptake was insensitive to NLRP3 inhibition by MCC950 (Figure 2A and 2B). In parallel sets of stimulations, MCC950 prevented LDH release and PI uptake that was induced by the treatment of LPS-primed cells with the NLRP3 agonist nigericin (Figure 2C and 2D). These collective results establish an experimental system to induce GSDMD pore forming activity directly, thus enabling the identification of factors that specifically mediate these events.

**Figure 2:**
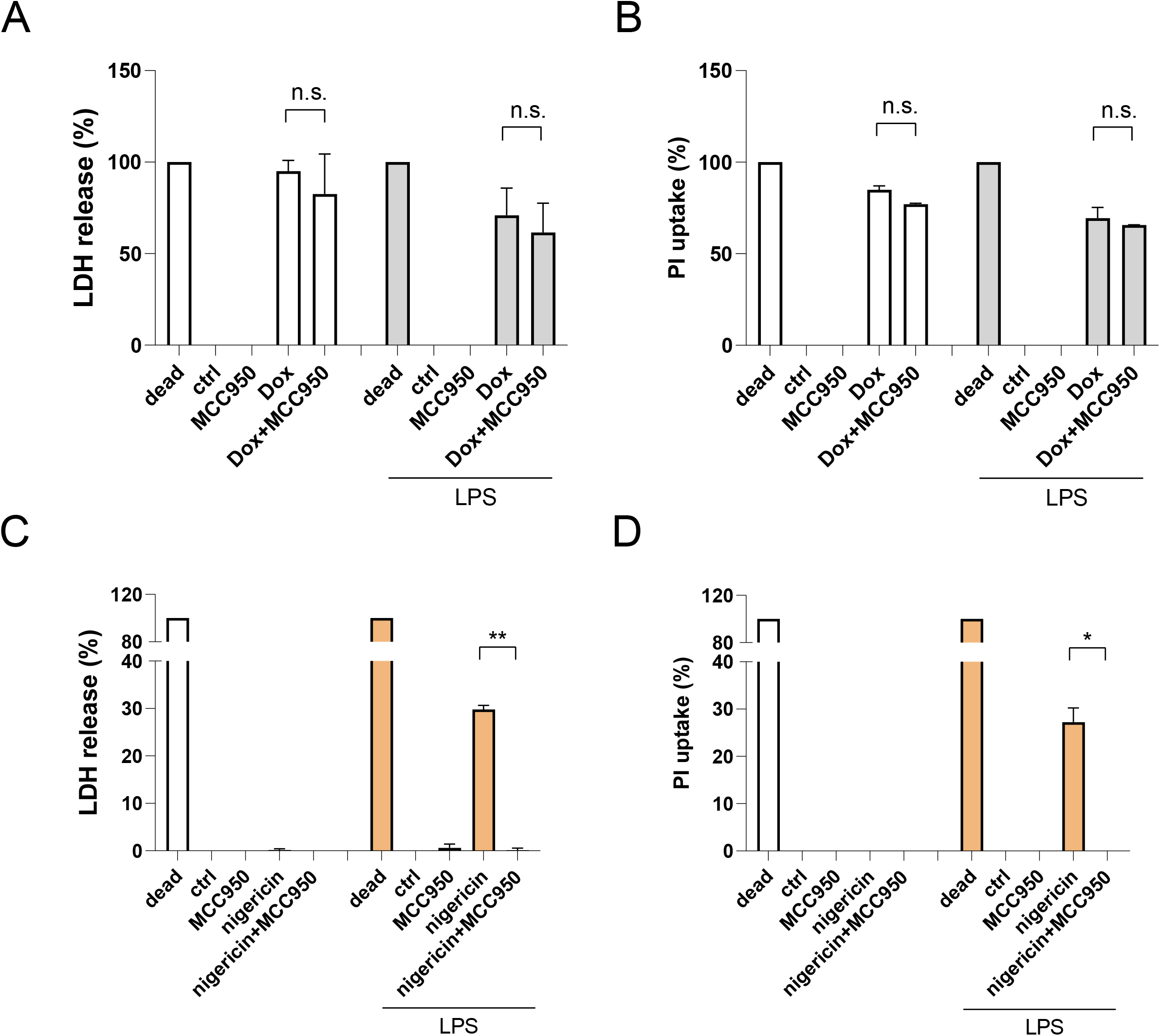
Inhibition of NLRP3 inflammasome does not affect pore formation and pyroptosis induced by NT-GSDMD. (A) End point analysis of PI uptake by plate reader to measure bulk membrane permeability in populations of doxycycline induced (0.5 μg/ml) NT-GSDMD-BFP clone 2 iBMDMs with or without MCC950 pretreatment (5 μM for 30 minutes) for 13 hours. (B) End point analysis of LDH release into cell free supernatants to measure bulk cellular lysis in populations of doxycycline induced (0.5 μg/ml) NT-GSDMD-BFP clone 2 iBMDMs with or without MCC950 pretreatment (5 μM for 30 minutes) for 13 hours. (C) End point analysis of PI uptake by plate reader to measure bulk membrane permeability in populations of LPS primed (100 ng/ml for 5 hours) and Nigericin stimulated (5 μM for 1 hour) WT iBMDMs with or without MCC950 pretreatment (5 μM for 30 minutes). (D) End point analysis of LDH release into cell free supernatants to measure bulk cellular lysis in populations of LPS primed (100 ng/ml for 5 hours) and Nigericin stimulated (5 μM for 1 hour) WT iBMDMs with or without MCC950 pretreatment (5 μM for 30 minutes). Data normalized to 100% lysis condition and unstimulated media controls. All quantification represents the mean and SEM of two independent experiments.

### A forward genetic screen identifies the Ragulator-Rag complex as factors necessary for GSDMD pore formation

To identify genes that regulate GSDMD pore formation, a genome-wide CRISPR-Cas9 screen was performed. Single cell clones of the iBMDMs encoding NT-GSDMD were first verified to produce similar amounts of Cas9 protein, as compared to their parental line (Figure 1B). We lentivirally transduced iBMDMs encoding NT-GSDMD with the Brie genome-wide single guide RNA (sgRNA) library (Doench et al., 2016). The Brie library contains expression constructs for sgRNA with coverage of 4 guides per gene in the murine genome and non-targeting controls, which is coupled to the expression of a blasticidin resistance gene for selection of productive transduction.

We optimized transduction, spinfection and viral titer to approximate one viral integration per infected cell, which translated to a target transduction efficiency of 30-50%. Transduced cells were selected with blasticidin and pooled after 6 days of expansion for downstream assays. Cells were left uninduced or treated with Dox to induce NT-GSDMD expression for 16 hours. Induced cells were then stained with PI to label cells containing functional GSDMD pores. The top 15% of BFP-positive cells, PI-negative cells were isolated by FACS. These cells have expressed the NT-GSDMD transgene (as identified by the BFP fusion tag) but have not formed pores in the plasma membrane. These cells were therefore predicted to contain mutations in genes that promote GSDMD pore formation.

Genomic DNA was isolated from the entire uninduced cellular cohort to serve as an input sample to account for bias or changes to sgRNA abundance in the population of cells used in our study. After PCR amplification of sgRNAs from genomic DNA, and subsequent next generation sequencing, the log-normalized sgRNA abundance from the input sample was subtracted from the log-normalized sgRNA abundance of BFP-positive survivors. Using hypergeometric analysis from the Broad Genomic Perturbation Platform, average p-values were generated and plotted against average log-fold change (LFC) at the gene level (Figure 3A). Gene ontology analysis of the rank list of genes from most positive LFC to most negative LFC identified functional annotations that were highly-enriched in our dataset. The most significant function was attributed to positive regulation of mTOR signaling (Figure 3B). Many of the genes with the highest positive LFC and most significant p-values were part of the Ragulator-Rag complex (Figure 3A and 3C). Highlighted hits are indicated in red on the volcano plot (Figure 3A) and on a schematized cryo-EM structure of this protein complex (Figure 3C) (Shen et al., 2019). Specifically, we identified the GTPases RagA and RagC and their functionally-related GTPase activating protein (GAP) FLCN, as well as the major components of the Ragulator complex—Lamtor-1, −2, −3, −4. These proteins are best known for their metabolic activities (Liu and Sabatini, 2020), but a role in GSDMD activities is unprecedented. We focused our subsequent studies on validating and understanding their functions in pyroptosis.

**Figure 3:**
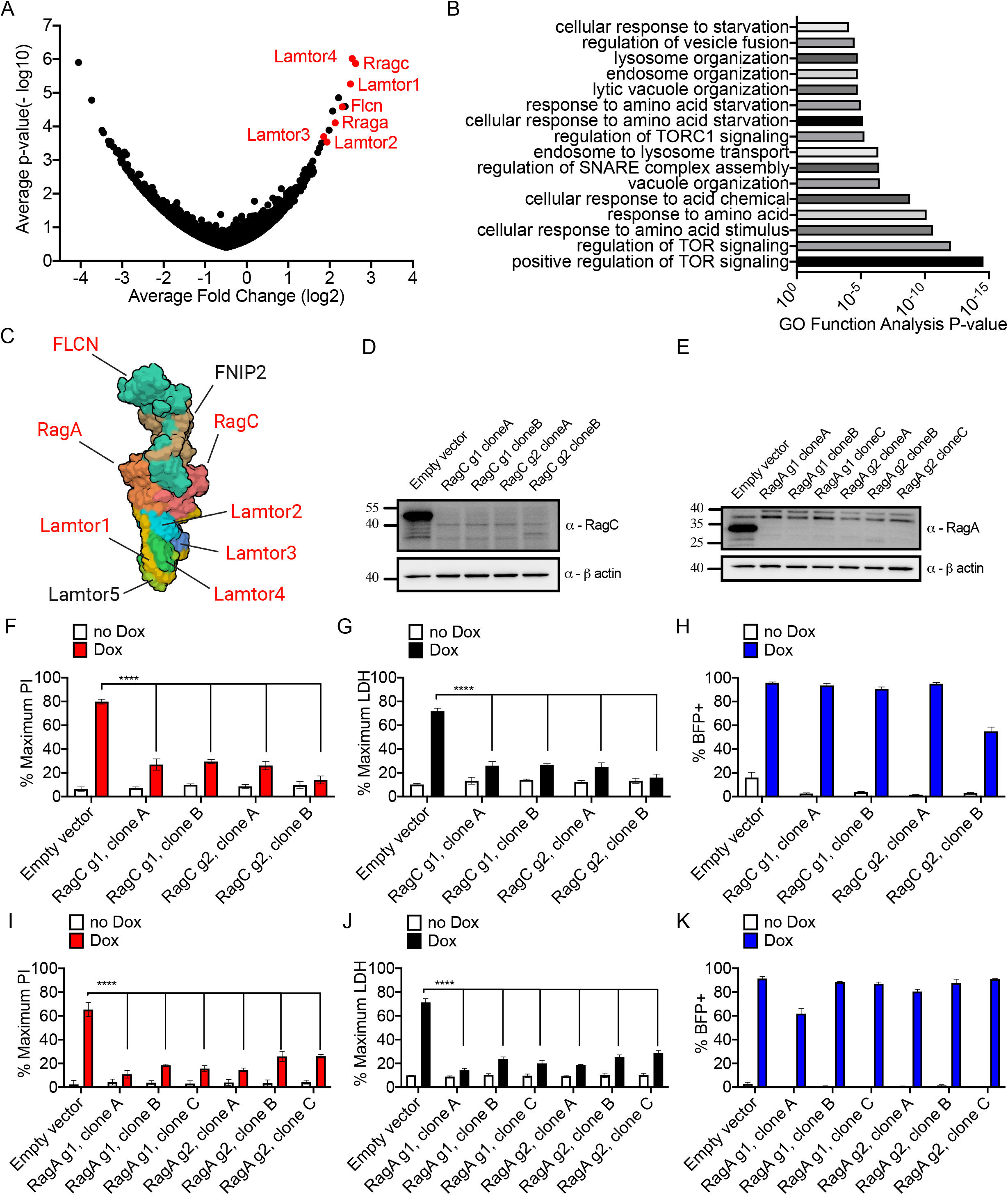
Analysis of survivor cells sgRNA enrichment and validation of screen hits. (A) Hypergeometric analysis of log normalized guide abundance of survivor cells subtracted by the log normalized abundance of input cells plotted as gene level average p-value versus gene level average log fold change. GSDMD pore formation potential positive regulators trend to the right of this plot and potential negative regulators trend to the left of this plot. Hit highlights and labeled in red are members of the Ragulator-Rag complex. (B) Gene ontology functional annotation enrichment analysis based on the complete hit list ranked from most positive LFC to more negative LFC using GOrilla web analysis tool. (C) Recent cryo-EM structure of the Ragulator-Rag complex cartoon schematic with top hits identified from CRISPR screen highlighted in red. (D) Western blot of RagC protein ablation comparing empty vector transduced or 2 independent clones and 2 independent sgRNA guides targeting RagC. (E) Western blot of RagA protein ablation comparing empty vector transduced or 3 independent clones and 2 independent sgRNA guides targeting RagA. (F, I) PI uptake endpoint plate reader membrane permeability of live cells left uninduced or doxycycline induced (2 μg/ml) for 16 hours comparing empty vector transduced or (F) 2 independent clones and 2 independent sgRNA guides targeting RagC or (I) 3 independent clones and 2 independent sgRNA targeting RagA. (G, J) LDH release into cell-free supernatants enzymatic endpoint plate reader cellular lysis assay from cells left uninduced or doxycycline induced (2 μg/ml) for 16 hours comparing empty vector transduced or (G) 2 independent clones and 2 independent sgRNA guides targeting RagC or (J) 3 independent clones and 2 independent sgRNA targeting RagA. (H, K) Frequency of BFP+ cells endpoint NT-GSDMD induction assay of live cells left uninduced or doxycycline induced (2 μg/ml) for 16 hours comparing empty vector transduced or (H) 2 independent clones and 2 independent sgRNA guides targeting RagC or (K) 3 independent clones and 2 independent sgRNA targeting RagA. All quantification represents the mean and SEM of three independent experiments.

To validate our top screen hits, we targeted specific components of the Ragulator-Rag complex for genetic ablation. Using two independent sgRNAs and two independent single cell clones from these sgRNAs, we ablated protein production of RagC, as compared to empty vector transduced cells (Figure 3D). Similarly, using two independent sgRNAs and three independent single cell clones from these sgRNAs, we ablated protein production of RagA, as compared to empty vector transduced cells (Figure 3E). After induction of NT-GSDMD, RagA-or RagC-deficient clones displayed defects in pore formation, as assessed by PI staining (Figure 3F and 3I). After induction of NT-GSDMD, RagA-or RagC-deficient clones also displayed defects in cell lysis, as assessed by LDH release (Figure 3G and 3J). Importantly, the percentage of BFP-positive cells was comparable between Rag-deficient cells and cells containing an empty vector control (Figure 3H and 3K). The comparable BFP expression observed indicates that the defects in pore formation associated with Rag-deficiency occur despite these cells expressing the NT-GSDMD transgene. Live cell imaging verified these results, as empty vector control cells displayed high frequencies of PI uptake and disintegration of cellular morphology after induction of NT-GSDMD (Video S1). In contrast, a low frequency of PI uptake and maintenance of cellular morphology was observed in two different lines of RagA-deficient iBMDMs (Video S2-S3). These collective data validate the importance of Ragulator-Rag in GSDMD-mediated pore formation and pyroptosis.

### RagA is not required for NT-GSDMD plasma membrane localization, but is necessary for pore formation and mitochondrial inactivation

Co-incident with the ability to form pores in the plasma membrane is the localization of NT-GSDMD to this location. Also co-incident with pore formation by GSDMD is the disruption of mitochondrial activities (de Vasconcelos et al., 2019). The combined effects of pore formation and mitochondrial disruption are considered necessary for a cell to ultimately lyse. In order to determine the mechanism by which Ragulator-Rag impacts NT-GSDMD activities, we examined the role of RagA in these events that occur in the pathway to pyroptosis. These efforts were focused on the behaviors of individual cells.

We first investigated whether the trafficking of NT-GSDMD to the plasma membrane is impacted by RagA deficiency after Dox-induction of NT-GSDMD expression. As expected, WT cells displayed NT-GSDMD at the plasma membrane and co-stained intensely with PI (Figure 4A). Within these cells, the plasma membrane displayed the expected ballooning morphology that is associated with GSDMD pore formation and pyroptosis. Consistent with our earlier assays (Figure 3I), we observed that RagA-deficient cells prevented the NT-GSDMD mediated uptake of PI (Figure 4A and 4B). Also in contrast to WT counterparts, RagA-deficient cells did not display a ballooning plasma membrane morphology (Figure 4A). Notably, despite the inability to permit PI uptake or balloon the plasma membrane, RagA-deficient cells displayed NT-GSDMD at the cell surface (Figure 4A). These data demonstrate that RagA does not control NT-GSDMD plasma membrane localization, but is rather required for pore forming activity after recruitment to this location.

**Figure 4:**
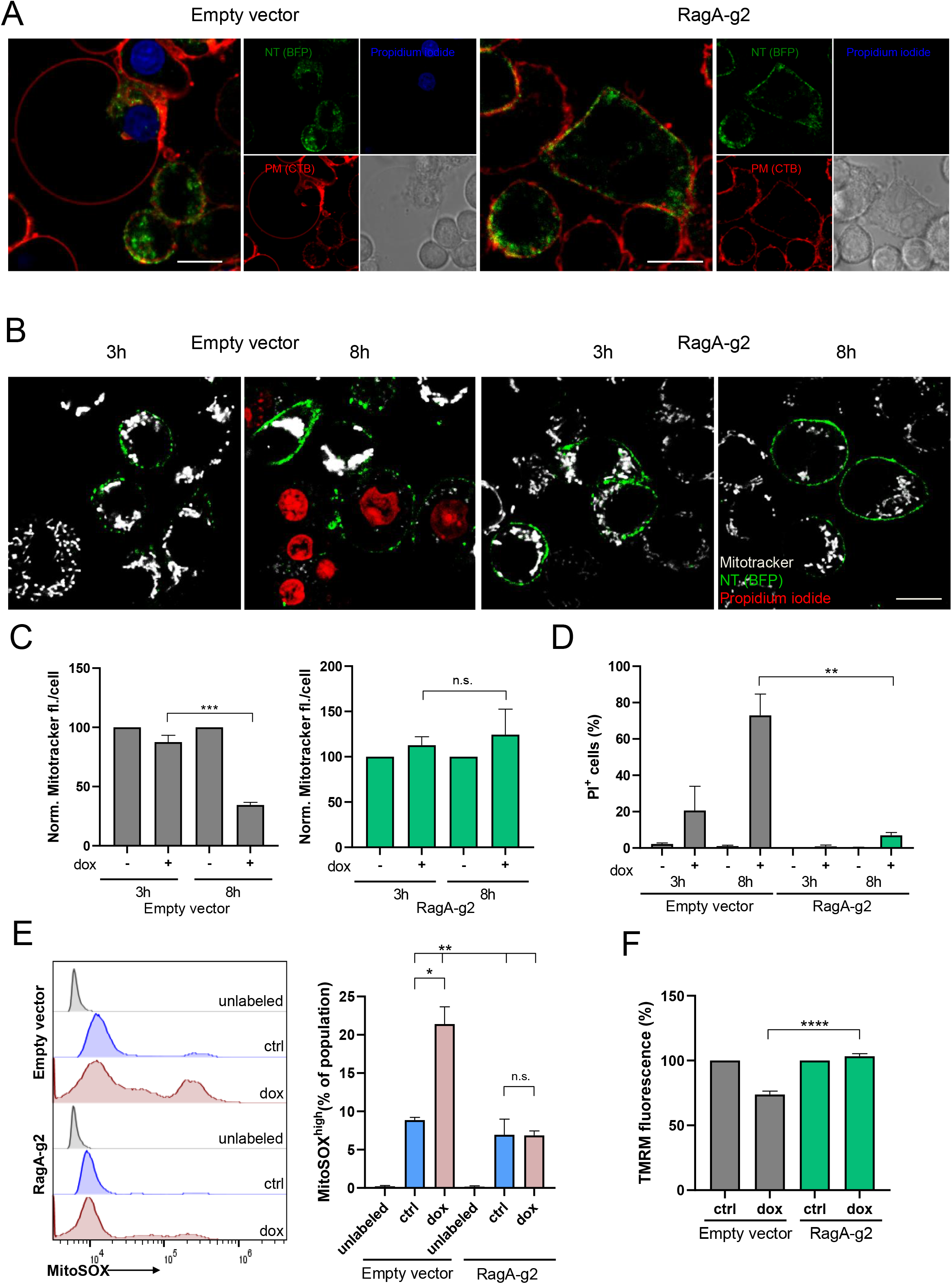
Rag deficiency does not affect localization of NT-GSDMD to the plasma membrane, but compromises functional pore formation and mitochondrial inactivation. To assess localization and mitochondrial health, empty vector transduced and RagA-g2 transduced clone C NT-GSDMD expressing cells were used. (A) Confocal microscopy of NT-GSDMD (green) expression was followed at 3 and 8 hours post doxycycline induction (0.5 μg/ml) in empty vector transduced or a clone of sgRNA targeting RagA transduced cells. Plasma membrane was labeled with Cholera toxin B-AF647 (cyan) and pore formation was followed by PI (red). Scale bar indicates 10 μm. (B) Live cell confocal microscopy of mitochondria labeled with Mitotracker Deep Red stain (cyan), nuclei of porous cells with PI (red) and NT-GSDMD expression (green) in empty vector transduced or a clone of sgRNA targeting RagA transduced cells. Scale bar indicates 10 μm. (C) Quantification of Mitotracker Deep Red high cells after doxycycline induction in empty vector transduced or a clone of sgRNA targeting RagA transduced cells. To quantify PI-positive cells (C) and cells enriched in Mitotracker Deep red stain (D) 3-5 246×246 μm^2^ were analyzed per experimental point. Mitotracker fluorescence was normalized to 100% for cells where doxycycline was not added. Mean and the SEM of three independent experiments are shown. Unpaired two-tailed t-test was used for analysis. (D) Quantification of PI-positive cells after doxycycline induction in empty vector transduced or a clone of sgRNA targeting RagA transduced cells. Unpaired two-tailed t-test was used for analysis. (E) Quantification of mitochondrial ROS induction as followed by MitoSOX stain at 8 hours post doxycycline addition in empty vector transduced or a clone of sgRNA targeting RagA transduced cells. Representative histograms are shown on the left and mean and SEM of MitoSOX-enriched fraction of intact cells from three independent experiments are shown on the right. Two-way ANOVA with Tukey multiple comparison test correction was used for analysis. (F) TMRM fluorescence at 8 hours post doxycycline induction in empty vector transduced or a clone C of sgRNA2 targeting RagA transduced cells followed by plate reader, normalized to noninduced controls. Unpaired two-tailed t-test was used for analysis. Mean and SEM of five independent experiments is shown.

Ninjurin-1 (NINJ1), a cell surface adhesion protein, was recently discovered to control pyroptosis at a late stage (Kayagaki et al., 2020). NINJ1 is required to induce plasma membrane rupture downstream of GSDMD pore formation. Ninj1-deficient cells energetically die upon inflammasome stimulation, as judged by the loss of mitochondrial activities, yet persist in ballooned morphology. As PI uptake was inhibited and ballooned morphology was not observed in RagA-deficient cells, in contrast to WT cells (Figure 4A), we examined the role of RagA in mitochondrial activities. We first followed mitochondrial integrity by confocal microscopy using Mitotracker Deep Red dye (Figure 4B, Supplementary Figure 1A). Dox-induced NT-GSDMD expression in WT iBMDMs coincided with a decrease in Mitotracker staining, whereas RagA-deficient cells maintained Mitotracker fluorescence upon NT-GSDMD expression (Figure 4B, Supplementary Figure 1A and 1B). This finding is inversely correlated with the genetic trends observed upon PI staining (Figure 4B and 4D, Supplementary Figure 1A-C). Thus, an increase in PI staining correlates with a loss of Mitotracker fluorescence, both of which occur in WT cells upon NT-GSDMD expression (Figure 4B-D, Supplementary Figure 1A-C). As an extension of our analysis of mitochondrial health, we assayed mitochondrial reactive oxygen species (ROS), which typically identifies stressed or damaged mitochondria (Ip et al., 2017). We found that mitochondrial ROS staining increased greatly in WT cells upon expression of NT-GSDMD (Figure 4E). In contrast, NT-GSDMD did not stimulate mitochondrial ROS in RagA-deficient cells (Figure 4E). Finally, RagA-deficient iBMDMs maintained mitochondrial membrane potential upon NT-GSDMD induction, as assessed using the potentiometric mitochondrial dye TMRM (Figure 4F and Supplementary Figure 1D). WT cells, in contrast, experienced a decrease in membrane potential upon NT-GSDMD induction (Figure 4F and Supplementary Figure 1D).

Thus, while the localization of NT-GSDMD to the plasma membrane is not facilitated by Ragulator-Rag, the formation of functional pores at the plasma membrane is abrogated in RagA-deficient cells. Moreover, RagA-deficient cells are protected from indications of mitochondrial dysfunction that are associated with NT-GSDMD pore formation. These data together position the actions of Ragulator-Rag downstream of GSDMD cleavage and plasma membrane localization, but upstream of NINJ1-induced plasma membrane rupture.

### mTORC1, but not mTORC2, is required for endogenous GSDMD pore forming activity and pyroptosis

Two major questions emerge from the genetic screen we have performed. What is the function of the Ragulator-Rag complex in GSDMD activity, and do our findings extend to the natural processes of GSDMD pore formation and pyroptosis? To address these questions, we considered the known functions of the Ragulator-Rag complex. In the metabolic mTOR pathways, two protein complexes operate that are known as mTORC1 and mTORC2 (Liu and Sabatini, 2020). The former of these is controlled by the Ragulator-Rag complex in response to amino acid availability (Sancak et al., 2010). To determine the relative roles of mTORC1 and mTORC2 in pyroptosis, we examined the function of the mTORC1 component Raptor and the mTORC2 component Rictor in iBMDMs. iBMDMs were generated from mice encoding CRE recombinase whose expression is driven by the LysM promoter (hereafter referred to as LysM-CRE iBMDMs). Similarly, iBMDMs were derived from mice dually encoding LysM-CRE and homozygous floxed alleles of Raptor or Rictor, respectively (hereafter referred to as Raptor-deficient or Rictor-deficient iBMDMs). As the LysM gene is activated in the macrophage lineage during differentiation of BMDM from bone marrow, differentiation of the precursor primary cells was sufficient to cause protein ablation in LysM-CRE expressing Raptor flox/flox and LysM-CRE expressing Rictor flox/flox iBMDMs (Supplementary Figure 2A). These cells were then subjected to stimulations that activate the natural pathways leading to cleavage and pore formation by endogenous GSDMD.

We first focused on GSDMD pore forming activity that is stimulated upon detection of cytosolic LPS by caspase-11. LysM-CRE WT iBMDMs were primed with LPS and subsequently electroporated with PBS or LPS to activate caspase-11. Under these conditions, we observed robust induction of pyroptosis from LysM-CRE WT iBMDMs, as evidenced by PI incorporation and LDH release (Figure 5A and 5B). Co-incident with pyroptosis was the release of IL-1β into the extracellular space (Figure 5C). In contrast to these behaviors of LysM-CRE WT iBMDMs, Raptor-deficient cells were defective for pore formation, lysis and IL-1β release (Figure 5A-C). Rictor-deficient cells were not defective for PI staining or LDH release upon LPS-priming and electroporation (Figure 5A and 5B). Live cell imaging confirmed these findings, as most LysM-CRE WT iBMDMs (Video S4) and Rictor-deficient iBMDMs (Video S5) experienced PI uptake and disintegration of cellular morphology after LPS electroporation. In contrast, fewer Raptor-deficient cells converted to PI-positivity after LPS electroporation (Video S6). These data indicate that Raptor, but not Rictor, is required for LPS-induced pyroptosis. Interestingly, whereas Raptor-deficient cells released less IL-1β than WT counterpart cells, Rictor-deficient cells released a greater amount of IL-1β than WT counterparts (Figure 5C). Thus, when comparing the two major downstream effectors of the mTOR pathway, mTORC1 and mTORC2 likely have disparate (and possibly opposing) functions in the pathways to pyroptosis.

**Figure 5:**
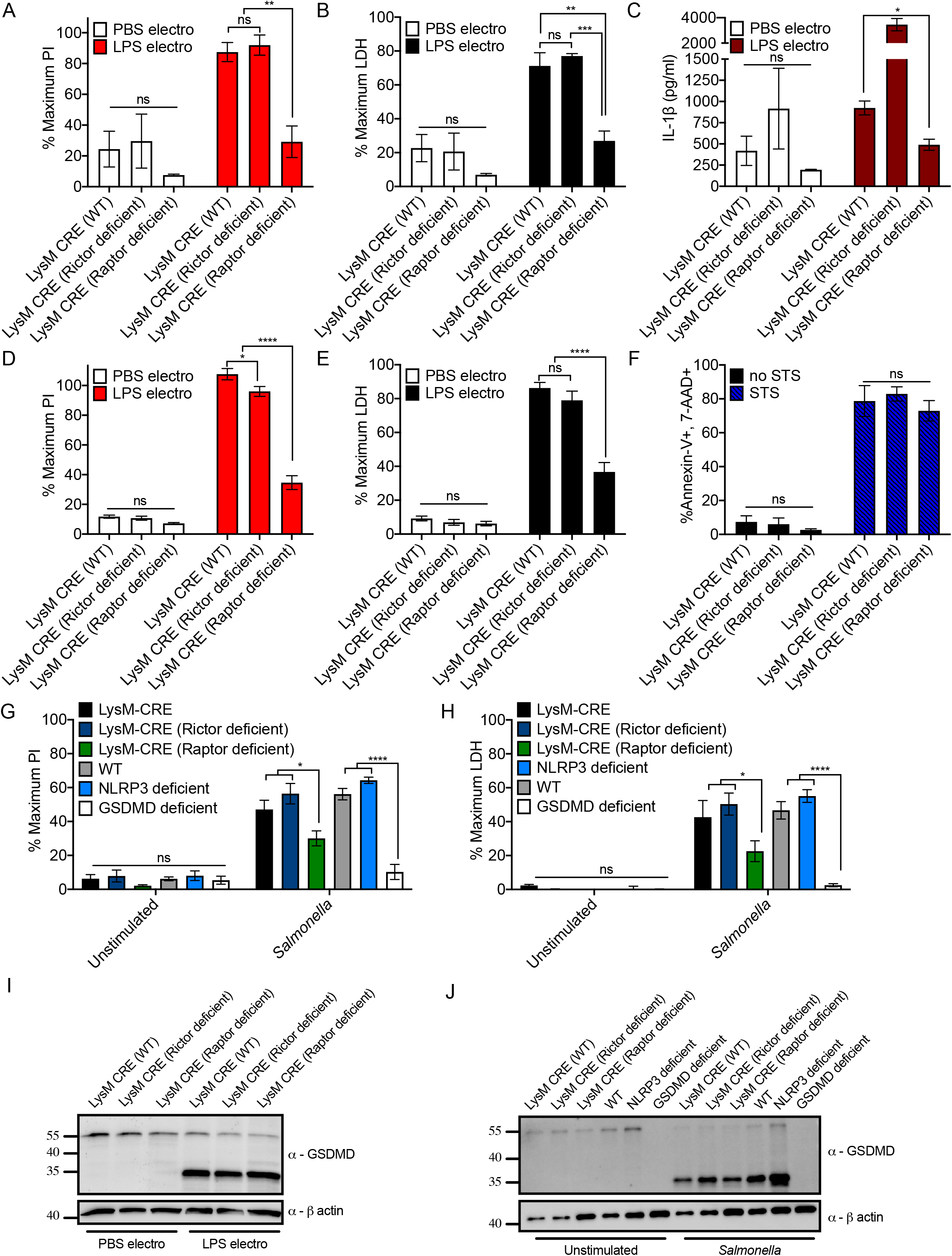
mTORC1 acts downstream of GSDMD cleavage to promote pore formation and pyroptosis in response to natural activation stimuli. (A) PI uptake endpoint plate reader membrane permeability of LysM-CRE WT, Rictor-deficient, and Raptor-deficient iBMDMs primed with LPS (1 μg/ml) for 3 hours then electroporated with PBS or 1 μg/million cells of LPS. Electroporated cells plated for 3 hours before analysis. (B) LDH release into cell-free supernatants was assessed from LysM-CRE WT, Rictor-deficient, and Raptor-deficient iBMDMs primed with LPS as in (A). Electroporated cells plated for 3 hours before analysis. (C) IL-1β ELISA measurements of cell-free culture supernatants from LysM-CRE WT, Rictor-deficient, and Raptor-deficient iBMDMs primed with LPS as in (A). Electroporated cells plated for 3 hours before analysis. (D) PI uptake endpoint plate reader membrane permeability of LysM-CRE WT, Rictor-deficient, and Raptor-deficient iBMDMs primed with 1000U of recombinant IFNβ for 3 hours then electroporated with PBS or 1 μg/million cells of LPS as in (A). Electroporated cells plated for 3 hours before analysis. (E) LDH release into cell-free supernatants from LysM-CRE WT, Rictor-deficient, and Raptor-deficient iBMDMs primed with 1000U of recombinant IFNβ for 3 hours then electroporated with PBS or 1 μg/million cells of LPS as in (A). Electroporated cells plated for 3 hours before analysis. (F) FACS quantification of Annexin-V / 7-AAD double positive cells from LysM-CRE WT, Rictor-deficient, and Raptor-deficient iBMDMs after 8 hours of 1 μM Staurosporine (STS) treatment. (G) PI uptake endpoint plate reader membrane permeability of LysM-CRE WT, Rictor-deficient, Raptor-deficient, C57/BL6J WT, NLRP3-deficient, and GSDMD-deficient iBMDMs unstimulated or infected with *S.* Typhimurium at a multiplicity of infection of 10 for 1 hour. (H) LDH release into cell-free supernatants from LysM-CRE WT, Rictor-deficient, Raptor-deficient, C57/BL6J WT, NLRP3-deficient, and GSDMD-deficient iBMDMs unstimulated or infected with *S.* Typhimurium at a multiplicity of infection of 10 for 1 hour. (I) Western blot of GSDMD with combined cell and supernatant lysate from LysM-CRE WT, Rictor-deficient, and Raptor-deficient iBMDMs primed with 1000U of recombinant IFNβ for 3 hours and then electroporated with PBS or 1 μg/million cells of LPS. Electroporated cells plated for 3 hours before analysis. (J) Western blot of GSDMD with combined cell and supernatant lysate from LysM-CRE WT, Rictor-deficient, Raptor-deficient, C57/BL6J WT, NLRP3-deficient, and GSDMD-deficient iBMDMs unstimulated or infected with *S.* Typhimurium at a multiplicity of infection of 10 for 1 hour. All quantifications represent mean and SEM of 3 independent experiments.

To complement these studies, we replaced the LPS priming step with a priming step using recombinant type I interferon (IFN). As IL-1β is not upregulated by IFN signaling, we focused on assays for pore formation (PI staining) and lysis (LDH release). Similar to our findings in LPS-primed cells, LPS electroporation of IFNβ-primed LysM-CRE WT iBMDMs stimulated PI staining and LDH release, which was reduced in Raptor-deficient cells (Figure 5D and 5E). In contrast, Rictor-deficient cells displayed no defects in PI staining or LDH release upon LPS electroporation (Figure 5D and 5E). PBS electroporation of all IFNβ-primed of cells yielded low pyroptotic responses that were not specific to any genotype of cell examined. In contrast to the requirement of Raptor for pyroptosis induction, WT, Raptor- and Rictor-deficient cells similarly committed to death upon treatment with apoptosis-inducer staurosporine (STS) (Figure 5F).

GSDMD acts at the terminus of several inflammasome pathways. We reasoned that if the Ragulator-mTORC1 axis does control pyroptosis after GSDMD cleavage, then Raptor-deficient cells should be defective for pyroptosis even under conditions where multiple inflammasome pathways are activated. To test this prediction, we performed infections with the pathogen *Salmonella enterica* serovar Typhimurium. *S*. Typhimurium infections can activate several inflammasome pathways, including those stimulated by caspase-11, NLRP3 and NAIP-NLRC4 (Broz et al., 2010; Chen et al., 2014; Franchi et al., 2006; Man et al., 2014). These inflammasomes all converge on the cleavage of GSDMD as an effector mechanism to pyroptosis. We infected iBMDMs with *S*. Typhimurium and examined PI staining and LDH release. Within one hour of infection, robust membrane permeability and cytolysis occurred in LysM-CRE WT and in C57BL/6J WT iBMDMs (Figure 5G and 5H). Consistent with literature demonstrating that multiple inflammasome pathways can be activated by *S*. Typhimurium, NLRP3 deficiency did not protect iBMDMs from pyroptosis during infection, whereas GSDMD-deficient cells were defective for these responses (Figure 5G and 5H). Notably, Raptor-deficient cells display significantly lower PI staining and LDH release upon *S*. Typhimurium infection, as compared to Rictor-deficient or LysM-CRE WT cells (Figure 5G and 5H).

To determine the stage of the pyroptotic process Raptor is necessary to execute, we examined early and late molecular events. An early event in the pyroptotic process is the upregulation of caspase-11 induced by extracellular LPS, a process mediated by TLR4 and its downstream adaptor TRIF (Rathinam et al., 2012). IFNβ also upregulates caspase-11 expression. We found that the abundance of caspase-11 marginally differed when comparing LPS-or IFNβ-treated LysM-CRE WT iBMDMs to their Raptor- and Rictor-deficient counterparts (Supplementary Figure 2B and 2C). Thus, neither Raptor nor Rictor deficiency are associated with defects in cell priming, an observation consistent with prior work (Moon et al., 2015). Despite caspase-11 upregulation in all genotypes of cells, we noted that within unstimulated cells, caspase-11 abundance differed between these genotypes (Supplementary Figure 2B and 2C). The significance of this phenotype is unclear, as recent work suggests that (in addition to caspase-11 upregulation), unknown mechanisms are associated with poising cells for inflammasome activities (Brubaker et al., 2020). Thus, we sought to examine the question of cell priming in a functional manner. We reasoned that if there were any functional differences associated with Raptor or Rictor deficiency, in terms of receptor-proximal signaling events, then cleavage of GSDMD should be impacted. To address this possibility, we monitored cleavage of GSDMD after the stimulations described above. Notably, in contrast to other regulators of pyroptosis that have been identified through genetic analysis, Raptor-deficient cells were not defective for GSDMD cleavage when cells were primed with IFNβ and subsequently electroporated with LPS (Figure 5I). Raptor-deficient iBMDMs also displayed a similar extent of GSDMD cleavage as other genotypes after *S*. Typhimurium infection (Figure 5J). These data are consistent with the stage in the pyroptosis pathway Raptor would be expected to operate, as our forward genetic screen was designed to identify factors that act downstream of GSDMD cleavage. The Ragulator-Rag complex and its downstream effector mTORC1 therefore serves as a terminal control system that is necessary for GSDMD pore formation.

## Discussion

In this study, a genetic screen was designed to specifically interrogate a single stage in the pyroptosis pathway. This approach was distinct from other forward genetic screens, where natural upstream stimuli were used. By focusing on a single (and poorly characterized stage), we bypassed the possibility of identifying previously-defined genes that are necessary for pyroptosis, such as regulators of cell priming or inflammasome assembly (Kayagaki et al., 2019; Kayagaki et al., 2015; Napier et al., 2016; Shi et al., 2015). This approach likely explains our ability to enrich for genes that act downstream of priming and inflammasome activities, and even act downstream of GSDMD cleavage. Moreover, our screen employed an internal control (the BFP tag on the GSDMD transgene) to ensure the continued expression of the pore-forming molecule throughout our assays. This quality-control strategy likely explains why we did not identify any regulators of transcription and translation in our dataset. Thus, we were able to focus on potential regulators of a process that was considered to be poorly regulated, namely the steps between cleavage of GSDMD and plasma membrane pore formation. Current models of pyroptosis invoke the process of membrane repair as a negative regulatory mechanism to limit GSDMD activities after this pore forming factor has been cleaved (Lieberman et al., 2019; Ruhl et al., 2018). It was unexpected that we could identify factors that would promote GSDMD pore forming activities post-cleavage, yet our forward genetic screen and subsequent analysis revealed Ragulator-Rag-mTORC1 as factors that act at this stage.

In the metabolic pathways, Ragulator-Rag operates on lysosomes to sense amino acid availability and induce appropriate mTOR-dependent cellular responses that depend on mTORC1 (Liu and Sabatini, 2020). While several localization-function questions arise from our findings, our mechanistic analysis has eliminated the possibility that GSDMD must interact with lysosome-associated factors prior to plasma membrane delivery. Indeed, we found that RagA is not required for NT-GSDMD localization to the plasma membrane. RagA is however necessary for functional pore formation and subsequent mitochondrial inactivation. As described earlier, Ninjurin-1 is a factor that also acts downstream of GSDMD cleavage to promote pyroptosis (Kayagaki et al., 2020). Our analysis allowed us to place Ragulator-Rag upstream of Ninjurin-1 activities, as the former is required for pore formation and Ninjurin-1 acts downstream of pore formation and mitochondrial inactivation to promote cytolysis. The links between GSDMD, Ragulator-Rag-mTORC1 and Ninjurin-1 that control pyroptosis are undefined and await further investigations.

A link between metabolism and cell death phenotypes has been investigated in many prior studies, including recent work demonstrating that the inflammasome components NLRP3 and GSDMD can be post-translationally modified by TCA cycle intermediates (Hooftman et al., 2020; Humphries et al., 2020). Moreover, metabolic dysfunction and mitochondrial damage have been considered to initiate death pathway signaling events upstream of apoptosome and inflammasome activities (Andersen and Kornbluth, 2013; Prochnicki and Latz, 2017). While it is possible that Ragulator-Rag-mTORC1 could also impact upstream events in the inflammasome pathways, our data reveal that a checkpoint exists after inflammasome activation and after cleavage of the normally lytic molecule GSDMD. The strongest evidence supporting this conclusion derives from the way we designed our genetic screen, which bypassed all upstream regulatory mechanisms that control GSDMD cleavage. Consequently, we can no longer conclude that formation of NT-GSDMD is equivalent to functional pore formation.

In summary, this study broadens the stages in the pyroptosis pathway that are regulated events to include activities downstream of GSDMD cleavage. These regulatory events are likely specific to activities within cells, as GSDMD has the intrinsic capacity to insert into liposomal membranes upon cleavage by caspase-11. Based on this idea, it stands to reason that additional cellular factors may exist that control pore formation, pyroptosis and other innate immune signaling pathways. The design of genetic screens that focus on single stages in a pathway (as opposed to the entire pathway) may allow for the complexity of cellular activities to be condensed, thereby revealing unexpected regulation of innate immune signal transduction.

## Acknowledgements

This work was supported by NIH grants AI133524, AI093589, AI116550 and P30DK34854 to J.C.K and AI138369 to C.L.E. J.C.K. holds an Investigators in the Pathogenesis of Infectious Disease Award from the Burroughs Wellcome Fund. C.L.E. is supported by the Harvard Herchel Smith and Landry Cancer Biology Fellowships. I.H.B. received a Fulbright visiting scholar grant. The work was supported by Slovenian Research Agency (project grant ARRS J3-1746 to I.H.-B. and core funding P4-0176). I.H.B. is a recipient of the ICGEB grant (CRP SVN18-01). We thank members of the Kagan lab, Prof. Roman Jerala and members of Department of Synthetic Biology and Immunology for helpful discussions. We would like to thank Mojca Benčina and Nataša Kopitar Jerala for methodological advices. Biorender was used to generate schematics in Figure 1A. We thank Ivan Zanoni for critical bone marrow reagents and John Doench for critical help in screen analysis.

## Author contributions

Conceptualization, C.L.E., I.H.B., J.C.K.; Methodology, C.L.E., I.H.B., J.C.K.; Investigation, C.L.E., I.H.B.; Visualization, C.L.E., I.H.B.; Writing – Original Draft, C.L.E.; Writing – Review & Editing, C.L.E., I.H.B., J.C.K.; Funding Acquisition, C.L.E., I.H.B., J.C.K.; Supervision, C.L.E., J.C.K.

## STAR METHODS

### Contact for reagent and resource sharing

Further information and requests for resources and reagents should be directed to and will be fulfilled by the Lead Contact, Jonathan C. Kagan.

### Experimental Procedures

#### Construct preparation

For inducible expression upon retroviral transduction, all constructs were inserted into pRETROX Tre3G plasmid (Clontech). Gene encoding mouse full-length GSDMD was ordered from Synobiological and tagBFP-encoding plasmid was from Evrogen. I105N point mutation was introduced by site-directed mutagenesis. mtagBFP-tagged GSDMD variants were prepared by overlap PCR using Phusion HF polymerase (Thermo Fisher Scientific). All GSDMD gene variants were restriction digested by BamHI and EcoRI restriction nucleases and inserted into pRETRO X Tre3G at BamHI/EcoRI sites.

For CRISPR/Cas9 targeting parental empty vector pXPR-054-Blast was digested with BsmbI restriction enzyme and the cut vector was purified by gel extraction. SgRNA corresponding DNA oligos were ordered from IDT and designed based on the targeting sequence of guides found in the Brie library. Forward oligo sequences had the sequence CACCG added 5’ to the sgRNA targeting sequence. Reverse complement oligo sequences had the sequence AAAC added to the 5’ end. Equal molar quantities of forward and reverse oligos were allowed to anneal to each other and were phosphorylated for efficient ligation to the purified and digested pXPR-054-Blast vector.

### Cell lines, Transfection, and Retroviral/Lentiviral Transduction

Bone marrow derived macrophages were differentiated from whole bone marrow derived from flushed tibia and femurs from C57BL/6J wild type, NLRP3 deficient, gasdermin D deficient, LysM CRE expressing, LysM CRE × flox/flox Raptor, LyM CRE × flox/flox Rictor, and Cas9 knock in mice. In brief, leg bones were surgically removed and cleaned of surrounding tissue. The bones were washed with sterile PBS and left in complete DMEM ((Gibco) with 10% FBS (Gibco) and Penicillin Streptomycin (Gibco) with added supplements of L-glutamine (Gibco) and sodium pyruvate (Gibco) during extraction. The ends of tibia and femur were cut and the bones were flushed with sterile PBS in a 10 ml syringe per bone and bone marrow from each individual mouse was pooled and pelleted at 400 × g for 5 minutes. ~10E6 bone marrow cells were used per non-TC coated 10 cm plate for differentiation. Differentiation was carried out in complete DMEM supplemented with 30% L929 supernatants containing M-CSF. J2 retrovirus secreting cells were passaged in complete DMEM and split every 2-3 days to prevent acidification and confluency. For immortalization viral supernatant production, flasks of J2 secreting cells were allowed to reach confluency and left to accumulate virus in their supernatants for 24 hours after reaching 100% confluency. Viral supernatants were spun at 400 × g to remove cell debris and passed through a 0.45 um syringe filter to ensure no cellular cross-contamination. Two rounds of immortalization viral supernatants were placed on differentiating macrophage cultures and supplemented with 30% L929 supernatants. Transduced cells were allowed to recover in fresh complete DMEM with 30% L929 supernatants then split 1:2 or 1:3 as immortalized macrophages began to form clusters / colonies in non-TC coated 10 cm plates. Cells were progressively starved of M-CSF by reducing the L929 supernatant supplementation concentration over approximately 1.5-2 months until cells were able to grow without M-CSF and be split 1:10 every 2-3 days. Immortalized bone marrow derived macrophages (iBMDMs) were cultured in complete DMEM without L929 supernatant supplementation. Cells were washed in PBS pH 7.4 containing 2.5 mM EDTA to detach cells for passage. Cells were passaged 1:10 every 2-3 days to prevent overcrowding and media acidification.

For generating stable expression of Tet3G in Cas9 KI iBMDMs, pantrophic retrovirus particles were produced by transfecting plasmids pCMV-VSV-G (1.5 μg) and pRETROX Tet3G (2.5 μg) (Clontech) in Platinum-GP (Cell Biolabs) packaging cell line with lipofectamine 2000 (10 μL). 3 days after transduction media was swapped for complete DMEM with 1.5 mg/ml G-418 for selection of productively transduced cells. For generating doxycycline inducible GSDMD variants in Cas9 KI Tet3G stable cells, amphotrophic retrovirus particles were produced by transfecting the respective pRETROX Tre3G plasmid (4 μg) into Gryphon Ampho (Alelle Biotechnology) packaging cell line with lipofectamine 2000 (10 μL). Media was swapped for complete DMEM with 10 μg/ml puromycin and 1.5 mg/ml G-418 3 days after transduction. These cells were further single cell cloned by limiting serial dilution to yield clonal populations for purposes of magnitude and synchronization of doxycycline induction responses.

For generating CRISPR/Cas9 targeted cell lines, lentivirus particles were produced by transfecting plasmids psPAX2, pCMV-VSV-G, and the transfer plasmid pXPR-054-Blast containing the sgRNA sequence of interest. Plasmids were transfected into HEK 293T cells in 10 cm dishes at a confluency of 50-70% with lipofectamine 2000 at a DNA to lipofectamine ratio of 1:3. Media was changed 8-12 hours post DNA transfection and viral supernatants were collected 24 hours post media change. Viral supernatants were spun at 400 × g to remove cellular debris then passed through a 0.45 μm PVDF filter via syringe. Viral supernatants were placed directly on growing Cas9 expressing iBMDMs and spinfected at 1250 × g for 45 minutes at 30 °C. These cell lines were selected with 10 μg/ml of Blasticidin to select for cells expressing sgRNA stably after transduction.

### Brie Library Creation and Screening Assay in Engineered Macrophages

Brie sgRNA only lentiviral library was obtained from the Broad Institute Genome Perturbation Platform with Blasticidin resistance marker. Viral tittering prior to bulk library creation optimized transduction conditions. Increasing amounts of Brie library virus stock (0-500μL) were added to 3E6 cells in prewarmed, complete DMEM with 5 μg/ml polybrene (in a constant final volume of 2 ml) in a 12-well plate. Spinfection transduction was performed by centrifugation at 1250 × g for 45 minutes at 30°C. After spinfection, 2 ml of complete DMEM was added and cells were incubated until the next day. Each condition was re-plated at lower density (1E5 cells/well of 6-well plate) in two wells to allow for ample room to grow during expansion and selection. In half of the wells, cells were selected with 4 μg/ml blasticidin for 4 days. At the end of selection and expansion, cell viability was measured to determine infection efficiency through a ratio of the number of blasticidin resistant cells and the number of non-selected cells for each condition. The target efficiency is 30-50% transduction efficiency by this assay to yield primarily 0.5-1 virus per productively transduced cell to ensure our downstream enrichment analysis could correlate guide abundance to a single perturbation (Doench et al., 2016).

Downstream screening assays should have 400 cells per single guide perturbation representation for statistical robustness during enrichment analysis (Doench et al., 2016). For our library creation, we utilized 9450 μl of stock virus on 189E6 engineered macrophages. After spinfection as described above, 2 ml of fresh complete DMEM was added to each well and cells were incubated for one day. 24 hours after spinfection, cells were detached with cold sterile PBS with 2.5 mM EDTA and replated into 175 cm^2^ TC filter flasks (12E6 starting cells / flask). After one more day, selection was initiated using 4 μg/ml of blasticidin in complete DMEM that also contained 10 μg/ml puromycin and 1.5 mg/ml G418 to ensure our synthetic pathway was still maintained and productively transduced cells expressing sgRNA would also be selected and expanded. After 6 additional days, surviving cells were pooled and plated into 10 cm TC-coated dishes. ~150E6 cells were induced with doxycycline to express the toxic NT-GSDMD molecule for 16 hours. Cells were detached with cold PBS with 2.5 mM EDTA and sequentially washed with MACS buffer (1-2% FBS, 2.5 mM EDTA in pH 7.4 sterile PBS) to get rid of cellular debris and focus sorting analysis on intact and surviving cells. Cells were stained with PI solution (3.3 μg/ml) and sorted on an Aria cell sorter (BD). Cells were assayed for BFP fluorescence in the PacBlue channel and PI florescence in the PE-A channel. The top 15% BFP+, PI-cells were sorted as cells that may contain interesting mutations in genes that promote GSDMD pore formation as they expressed the NT-GSDMD pore forming domain to high levels, but showed defective functional pore formation. Cells were sorted into PBS EDTA with 30% FBS to ensure efficient capture of cells/nuclei for downstream genomic DNA extraction. ~200E6 cells were collected that were left uninduced to serve as an input control to account for potential skewing of guide abundance in our population during the expansion and screening assay (for example for guides targeting known tumor suppressors that may outgrow in the population). Genomic DNA was isolated using Nucleospin Blood XL (Clontech) according to Broad Genome Perturbation Platform protocols. The gDNA was also subjected to PCR inhibitor clean up columns from Zymo Research (D6030) then diluted to the desired concentration and number of wells and sent on dry ice to the Broad Institute for PCR amplification and next generation Illumina Sequencing.

### Analysis of Screening Results

Log normalized sgRNA abundance was provided by the Broad GPP from input and BFP+/PI-survivor cells populations. Population bias during expansion and assay were corrected for by subtraction of input log normalized sgRNA abundance from log normalized sgRNA abundance from the BFP+/PI-survivor cell population. We used the Broad GPP web portal for further analysis: https://portals.broadinstitute.org/gpp/public/. A strict Chip file was used to map perturbations to sites in the genome, and the normalized data was subjected to hypergeometric analysis without replacement. In this method, the rank of sgRNAs is used to calculate gene p-values using the probability mass function of a hypergeometric distribution. P-values are generated by calculating the average −log_10_(p-value) in both directions and picking the more significant one. The top n% of sgRNAs per gene can be used to calculate the average p-value with this method. The average log-fold change per gene is also reported and this can be used to assess the magnitude of effect. Data displayed on the volcano plot requires a minimum of 3 out of 4 guides being present.

For gene ontology functional analysis of our screen results, the complete rank list based on most positive log-fold change (LFC) to most negative LFC was input into the GOrilla web tool: http://cbl-gorilla.cs.technion.ac.il/.

### Doxycycline Induction in Engineered Cell Lines

Cell lines of different genotypes were counted using a Luna 2 cell counter and plated at 1E5 cells / well in a black 96 well plate with optically clear bottoms in 200 μl of complete DMEM. After cells adhered (>6 hours resting in an incubator), media was replaced with warm complete DMEM or complete DMEM with doxycycline (0.5 μg/ml for initial characterization and 2 μg/ml for analysis of CRISPR/Cas9 edited clones). Cells were stained with diluted PI solution (final concentration 1:300) and lysis buffer or additional media were added to respective wells 30 minutes prior to a time point. The plate was spun at 400xg for 5 minutes to ensure all cells were at the bottom of the well prior to adding the plate to the plate reader. Population PI staining was assayed with a Tecan plate reader. The program settings for PI inclusion assay were bottom reading of fluorescence with an excitation wavelength of 530 nm with a bandwidth of 9 nm and emission wavelength of 617 nm with a bandwidth of 20 nm. Ideal gain was calculated from one of the positive lysed replicate wells to make the signal from that well be 70% of max detectable signal. The number of flashes was set to 25 with an integration time of 20 μs with no lag time or settle time. Supernatants from the spun down plate were taken for CyQuant LDH colormetric enzyme assay (Thermo) per the manufacturer’s instructions. Cells were lifted directly with MACS buffer (1-2% FBS, 2.5 mM EDTA in sterile pH 7.4 PBS) and assayed for frequency of PI+ cells and frequency of BFP+ cells on a Fortessa FACS machine (BD). Associated flow cytometry data was analyzed using FlowJo 10.0. For live cell microscopy, diluted PI (final concentration 1:300) was added to the wells right after doxycycline addition. The plate was allowed to equilibrate with PI for 30 minutes prior to imaging to allow for setting of exposure, gain, and focus settings based on lysed wells. The plate was placed in the Biotek Cytation with 5% CO2 and 37 C degree temperature incubation with bright field and red channel images taken every 30 minutes for 8 hours.

### Electroporation

Electroporation was conducted using the Neon transfection machine, pipette, tips, and associated commercial buffers (ThermoFisher Scientific). For experiments involving electroporation of LPS into cells, 1.2E6 cells were suspended in 120 μl of R buffer from the Neon transfection kit. Either 1.2 μl of PBS or with 1.2 μl of 1mg/ml LPS were added to the cell suspension prior to electroporation. Cells were drawn into the neon transfection pipette and electroporated with the parameters of Voltage=1400, pulse width=10ms, and pulse number=2. For protein isolation, 1E6 cells were dispensed into 500 μl of L-glutamine and sodium pyruvate supplemented, serum-free optiMEM to allow for analysis of whole lysate and supernatant combined through direct lysis with 5X SDS Laemmli buffer and TCEP reducing agent. For PI incorporation, LDH release, and supernatant collection for ELISA analysis of cytokines, the 100 μl containing 1E6 electroporated cells were dispensed into 900 μl of complete DMEM lacking antibiotics and 100 μl of this cell solution was dispensed into 6 wells for each condition in a black 96 well plate with optically clear plastic bottom. Of these wells, 3 wells served as internal lysis controls and 3 wells served as experimental conditions. Diluted PI solution (final concentration 1:300) was added to every well 30 minutes prior to the end time point. Lysis buffer or complete DMEM with no antibiotics were added to corresponding wells for PI and LDH normalization. Population PI staining was measured with a Tecan plate reader. For live cell microscopy, diluted PI (final concentration 1:300) was added to the wells right after plating. The plate was allowed to equilibrate with PI for 30 minutes prior to imaging to allow for setting of exposure, gain, and focus settings based on lysed wells. The plate was placed in the Biotek Cytation with 5% CO2 and 37 C degree temperature incubation with bright field and red channel images taken every 10 minutes for 1 hour.

### Characterization of Apoptosis

The PacBlue Annexin-V and 7-AAD apoptosis detection kit from Biolegend was used to characterize the apoptotic ability of LysM-CRE WT, Rictor deficient, and Raptor deficient iBMDMs in response to 1 μM of Staurosporin (STS) for 8 hours. The manufacturer’s staining protocol was followed. Cells were analyzed by a Fortessa FACS machine (BD). Double positive (Annexin-V+, 7-AAD+) cells were quantified by quartile gating comparing unstimulated control cells and STS stimulated experimental conditions using FlowJo.

### Bacterial Infection

*Salmonella enterica* serovar Typhimurium (SL1344) was streaked on Luria Broth (LB) agar plates with no antibiotic selection to yield single colonies. Single colonies were picked with a sterile tip and used to inoculate a liquid culture of normal LB in 2.5 ml of a 14 ml bacterial culture tube and left to shake at 37 C for 12 hours. Cultures were back-diluted 1:20 into 2.5 ml of high salt LB (LB + 0.3M NaCl) to encourage an invasive phenotype for 3 hours shaking at 37 C. Liquid culture was analyzed undiluted and 1:10 diluted in high salt LB compared to a blank of high salt LB alone to measure an accurate OD600. It was assumed that an OD600 was roughly 1E9 bacteria / ml for purposes of MOI calculations. 1 ml of liquid culture was spun at 10,000 × g in a microcentrifuge tube for 5 minutes at 4 C to pellet bacteria. This pellet was sequentially washed in sterile PBS pH 7.4 for 3 washes and spins. Calculated amounts of bacteria to yield an MOI of 10 were added to either L-glutamine / sodium pyruvate supplemented antibiotic free OptiMEM for protein isolation experiments or complete DMEM lacking antibiotics for plate reader experiments in a black 96 well plate with optically clear bottoms. Bacteria were spun down onto at 400 × g for 5 minutes to synchronize bacterial uptake by macrophage lines. 5X SDS laemmli buffer with TCEP reducing agent was added directly to protein isolation wells to yield a cellular lysate and low serum supernatant protein sample for western blot analysis. 30 minutes prior to end point, dilute PI solution (final concentration 1:300) was added to all wells in the 96 well plate. Lysis buffer or complete DMEM with no antibiotics were added to respective wells for PI and LDH normalization. At the end point time point of 1 hour, the plate was spun at 400 × g for 5 minutes to ensure all cells were at the bottom of the wells and PI fluorescence was read with a Tecan plate reader. Samples of supernatant were taken for LDH colorimetric enzyme assay.

### Confocal microscopy

To follow localization of tagBFP-tagged gasdermin variants cells were seeded into μ-slide (Ibidi). Cells were non-treated or treated with doxycycline (0.5 μg/ml) for the defined time. 30 minutes prior the end of incubation period propidium iodide (3.33 μg/ml) was added. Cholera toxin subunit B-Alexa Fluor 647 (2 μg/ml) was added as well in the experiment to follow plasma membrane localization. Cells were observed under a Leica TCS SP5 laser scanning microscope mounted on a Leica DMI 6000 CS inverted microscope (Leica Microsystems, Germany) with an HCX plan apo 63× (NA 1.4) oil immersion objective used for imaging. A 405-nm laser line of 20-mW diode laser was used for mtagBFP excitation, a 543-nm laser was used to follow propidium iodide, and a 633-nm laser was used for Cholera toxin subunit B-Alexa Fluor 647 excitation. To observe mitochondria, cells were followed at different time points after addition of doxycycline (0.5 μg/ml), Mitotracker Deep Red FM (30 nM) and propidium iodide (3.33 μg/ml).

For acquisition and image processing, Leica LAS AF software was used where threshold, brightness and contrast were adjusted. All images from the same experiment were processed in the same way using the same parameters. Mitotracker Deep Red FM fluorescence was calculated with particle analysis tool in ImageJ where also initial thresholding to reduce non-mitochondrial stain was performed. Cells from 3-5 246×246 μm^2^ frames per experimental point were counted with CellCounter plugin in ImageJ. All images from the same experiment were processed in the same way using the same parameters.

### Detection of mitochondrial ROS and mitochondrial potential

For detection of mitochondrial ROS, cells were detached after 8h-stimulation with doxycycline and incubated with MitoSOX (5 μM) for 20 minutes at 37°C in PBS/2% FBS. After washing step, samples were followed by flow cytometry using Cytek Aurora and analyzed using FlowJo.

Detection of change in mitochondrial potential was followed with TMRM (tetramethylrhodamine, methyl ester, perchlorate). Cells in 96-well black microtiter plates were stimulated with doxycycline (0.5 μg/ml) for 7h after which TMRM (final concentration 100 nM) was added for one hour. Medium was replaced with PBS/2% FBS and TMRM fluorescence was measured with Synergy Mx (Biotek) with 535 nm excitation and 600 nm emission settings. TMRM fluorescence was normalized to untreated control as 100%. To follow TMRM fluorescence using flow cytometry, cells were detached after 7.5 h-stimulation with doxycycline and incubated with TMRM (100 nM) for 30 minutes at 37°C in PBS/2% FBS. After washing step, samples were followed by flow cytometry using Cytek Aurora and analyzed using FlowJo. MFI of samples 8h after doxycycline addition was normalized to non-treated (100%) and unlabeled (0%) control.

### Antibodies and Reagents

LPS from E. coli, serotype 0111:B4 was supplied by Enzo (ALX-581-012-L002). Nigericin was purchased from Invivogen (tlrl-nig-5). Propidium iodide (PI) was purchased from Sigma (P4864-10ML). LDH activity was assayed using legacy Pierce™ LDH Cytotoxicity Assay Kit from ThermoFisher Scientific (88954) and replaced by CyQuant LDH Cytotoxicity Assay Kit from ThermoFisher Scientific according to the manufacturer’s protocol (C7026). Mouse IL-1β release was quantitatively measured from cell-free culture supernatants using the Invitrogen IL-1 beta Mouse Uncoated ELISA Kit (88-7013-88) according to the manufacturer’s protocol. For western blots, primary antibodies were used to detect proteins of interest including antibodies to tagBFP from Evrogen (AB233, 1:1000), Tet3G transactivator from Takara (631131, 1:1000), spCas9 from Cell Signaling Technologies (65832S, 1:1000) and from Abcam (ab191468, 1:1000), RagA from Cell Signaling Technologies (4357S, 1:1000), RagC from Cell Signaling Technologies (5466S, 1:1000), Raptor from Cell Signaling Technologies (2280S, 1:1000), Rictor from Cell Signaling Technologies (2114S, 1:1000), GSDMD from Abcam (ab209845, 1:1000), caspase-11 from Biolegend (647202, 1:500), and β actin from sigma (cat number, 1:5000) and Cell Signaling Technologies (cat number, 1:5000), Secondary antibodies to respective species with conjugated HRP were from Jackson ImmunoResearch or Abcam and used at 1:5000 (anti-mouse HRP 115-035-003, anti-rat HRP 112-035-003, anti-rabbit 111-035-003, anti-rabbit ab6721). Chemoluminescent signal substrate for HRP was SuperSignal West Pico (34577) and Femto Chemiluminescent Substrate (34095) from Thermo Scientific (cat number) and ECL from Amersham, GE Healthcare Life Sciences (RPN2232).

### Quantification and Statistical Analysis

Statistical significance for experiments with more than two groups was tested with twoway ANOVA with Tukey multiple comparison test correction. Unpaired two-tailed t-test was used for comparison of two groups. Adjusted p values were calculated with Prism 8.0 from GraphPad and the designation of * corresponds to p ≤ 0.05, ** corresponds to p ≤ 0.01, *** corresponds to p ≤ 0.001, and **** corresponds to a p ≤ 0.0001 in the figures. Data presented is representative of at least 3 independent experiments for western blots. Data presented as quantified bar graphs or time course analysis is the combined means of 3 independent experiments unless otherwise designated in figure legends. Data with error bars are represented as mean ± SEM.

## KEY RESOURCES TABLE

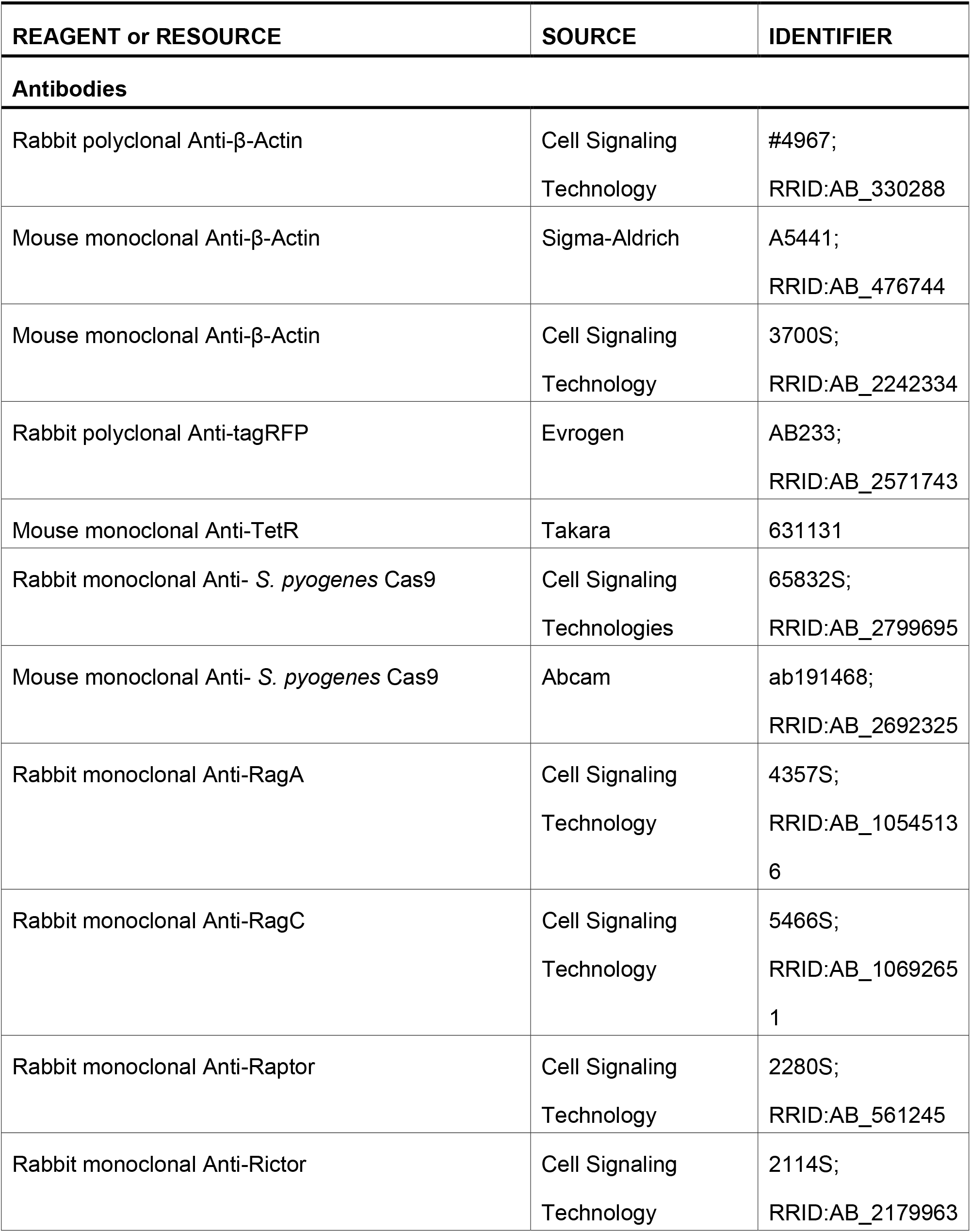
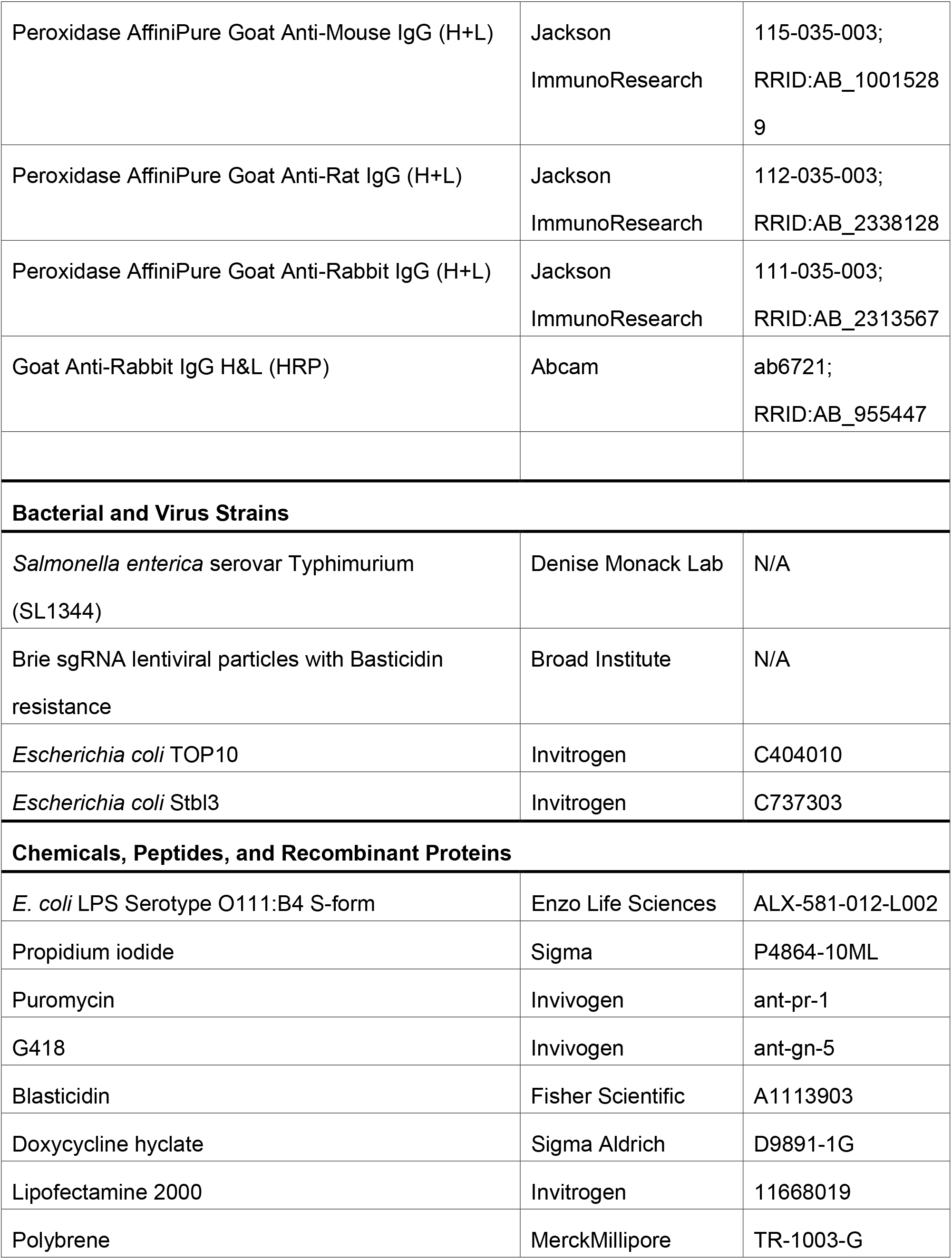
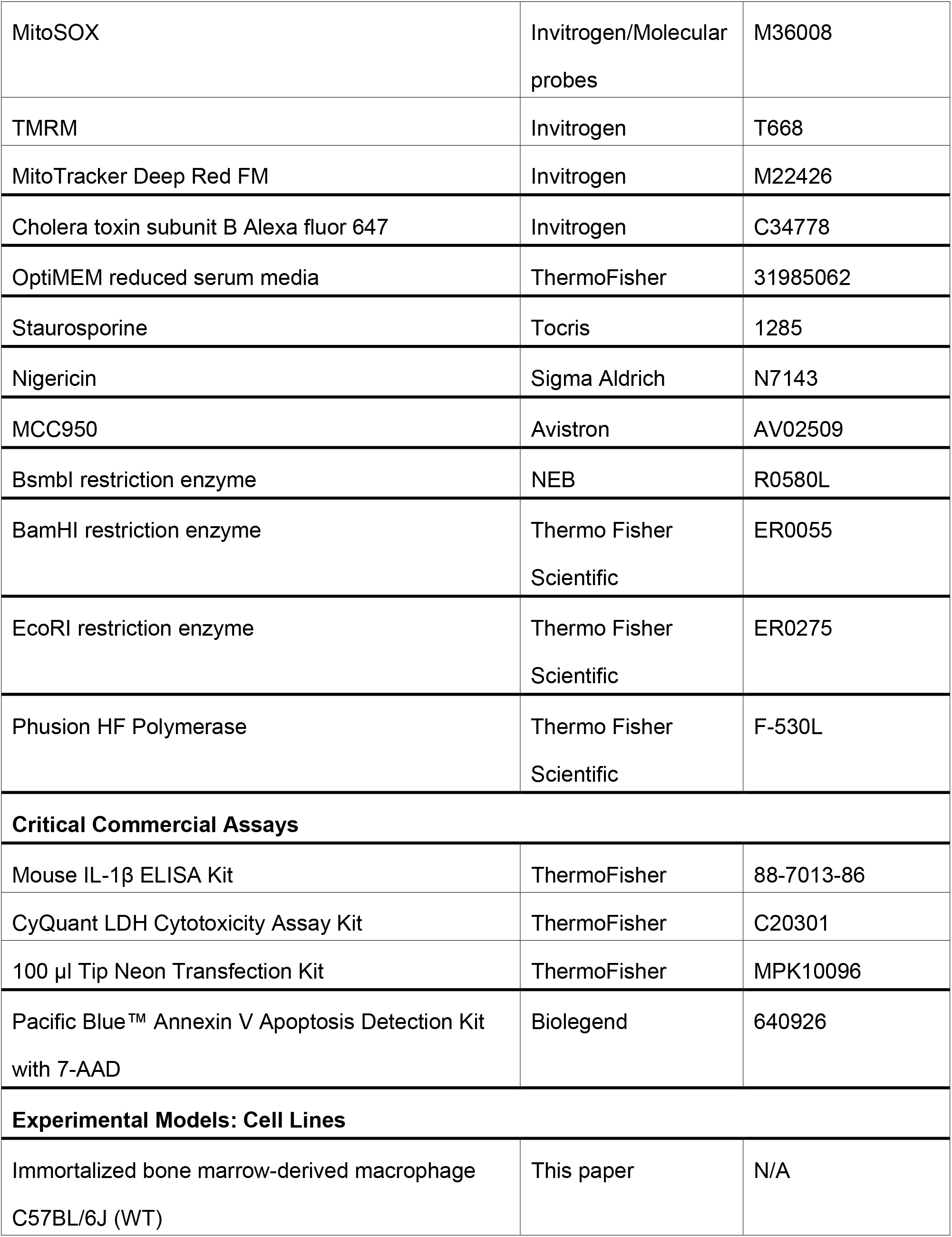
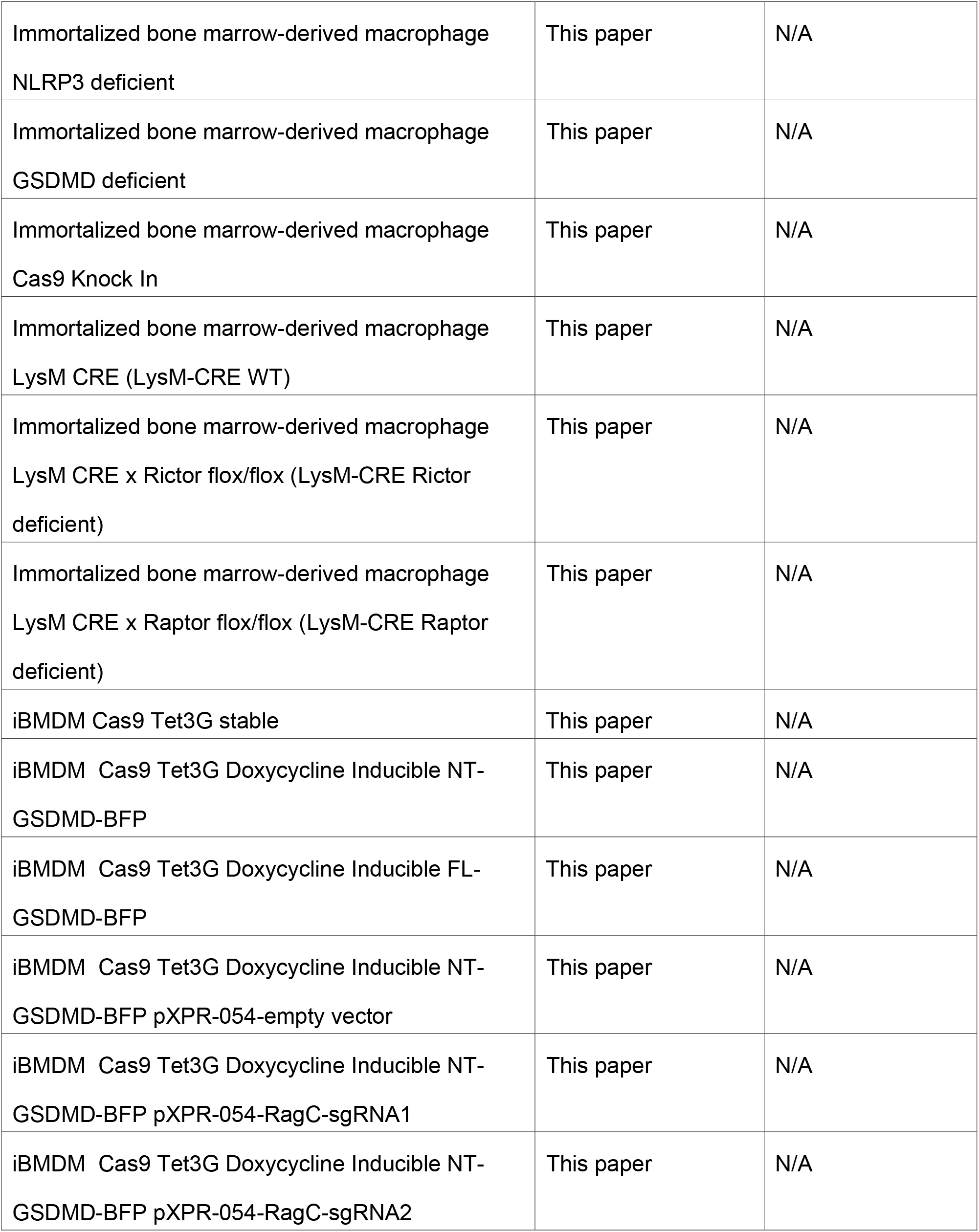
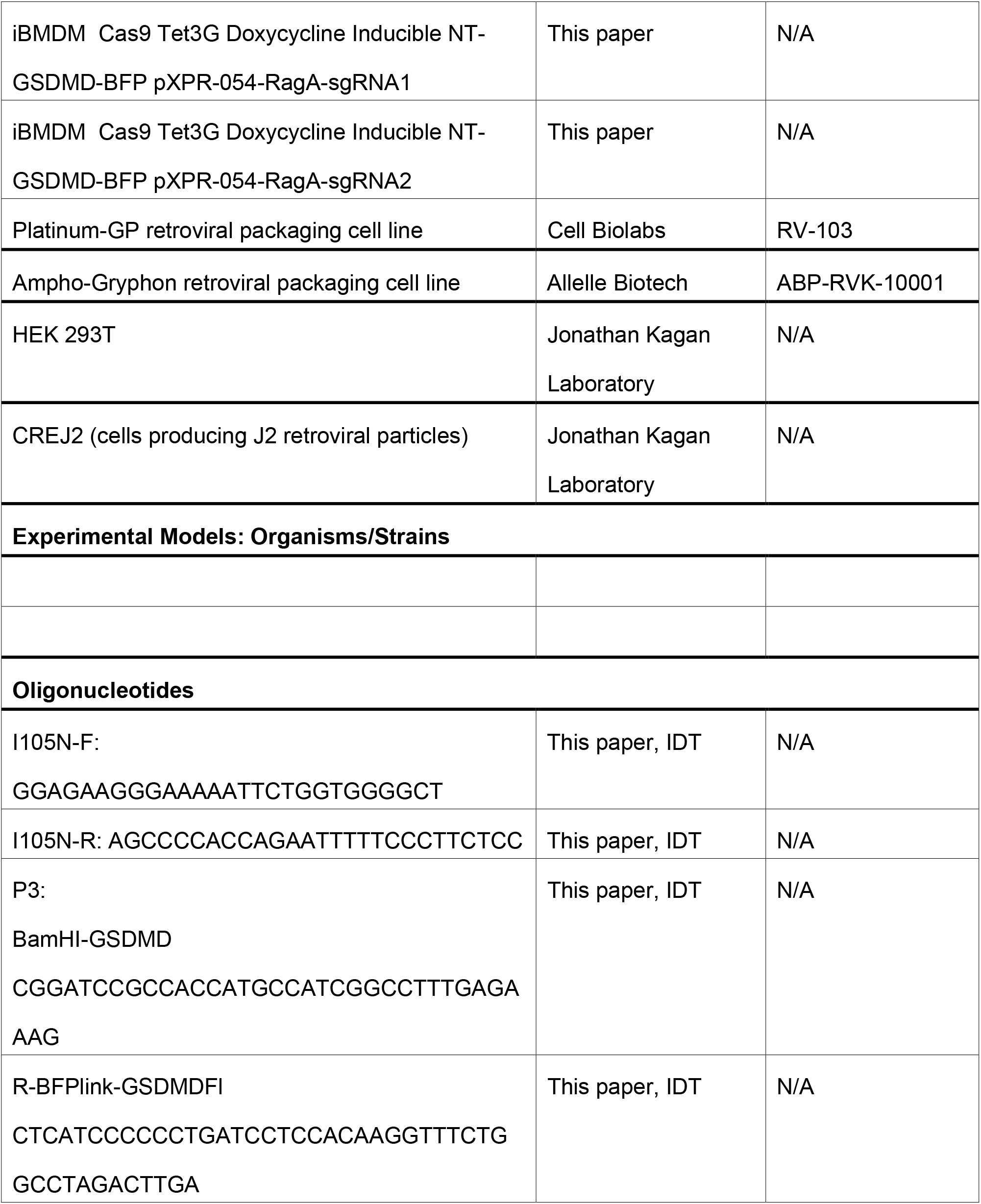
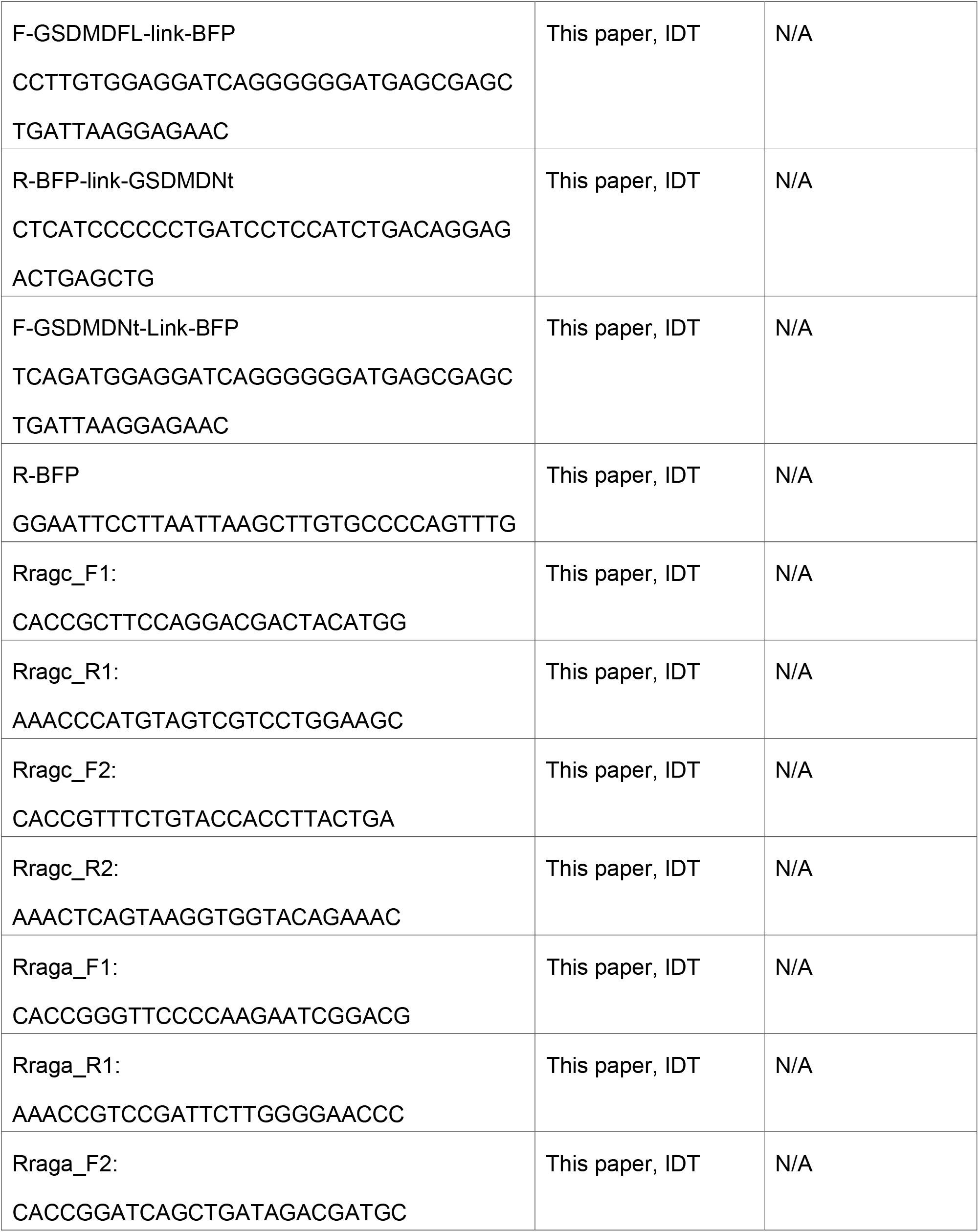
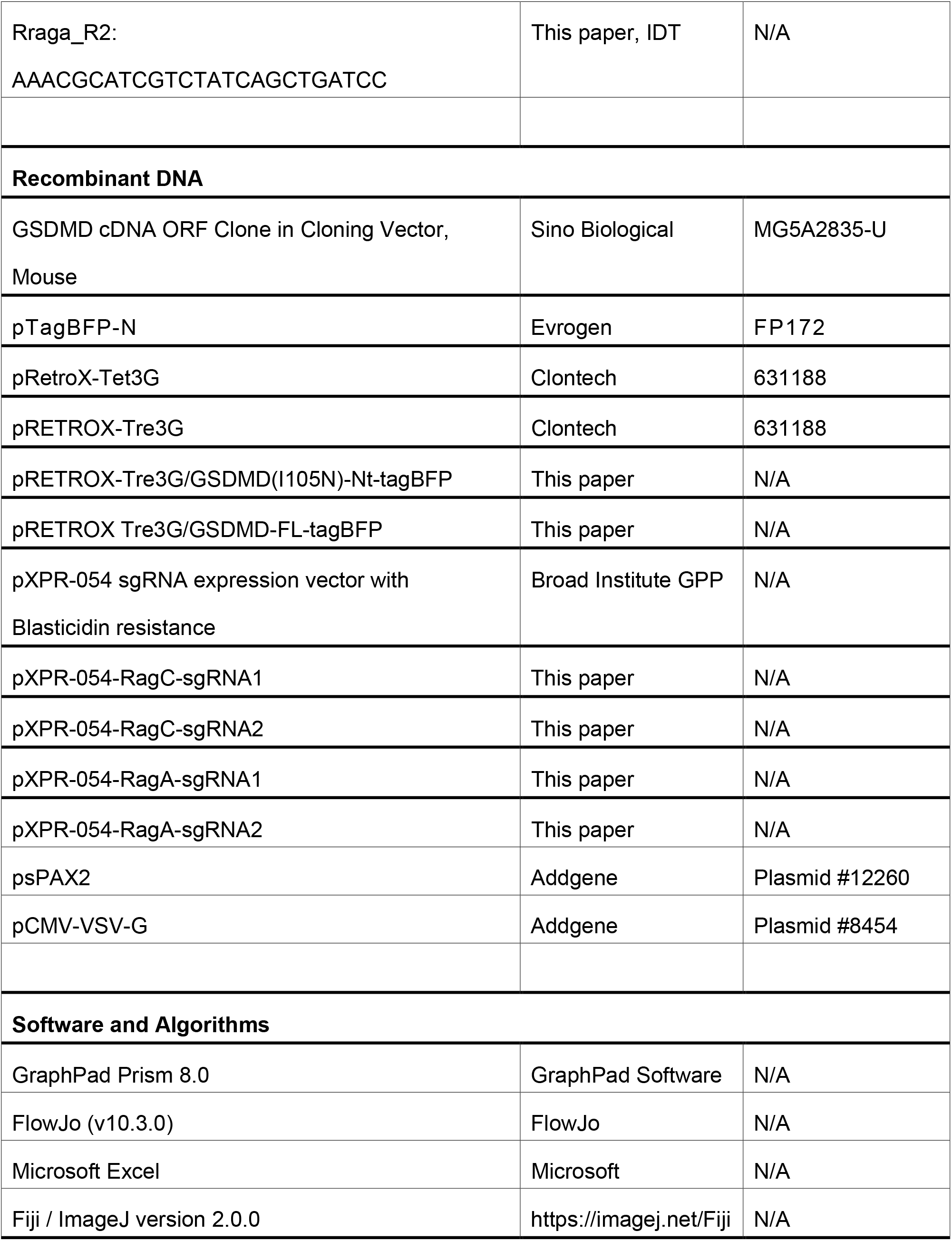
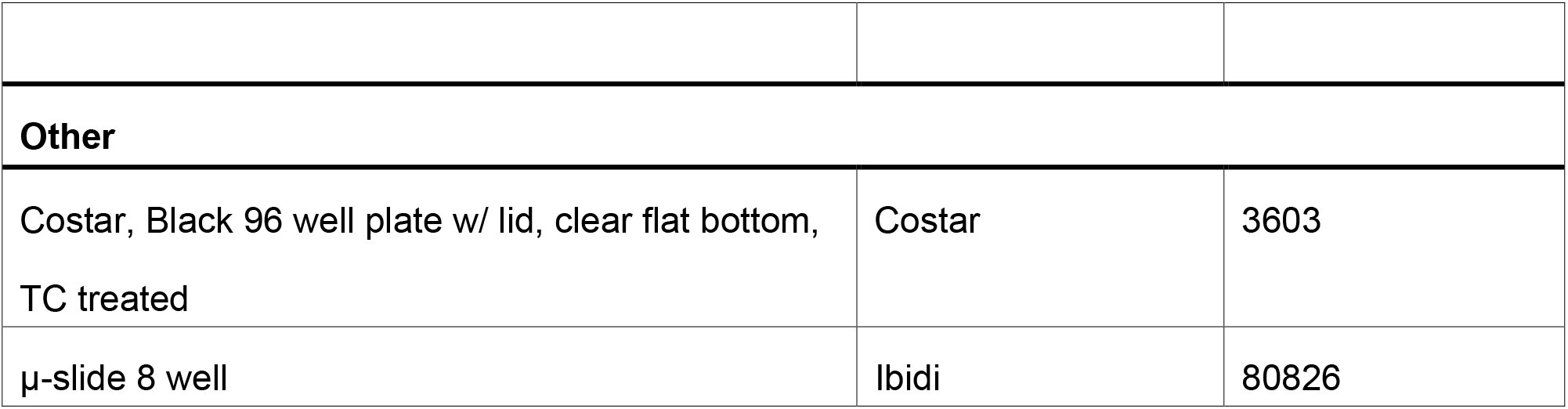

---

**Supplemental Figure 1:**
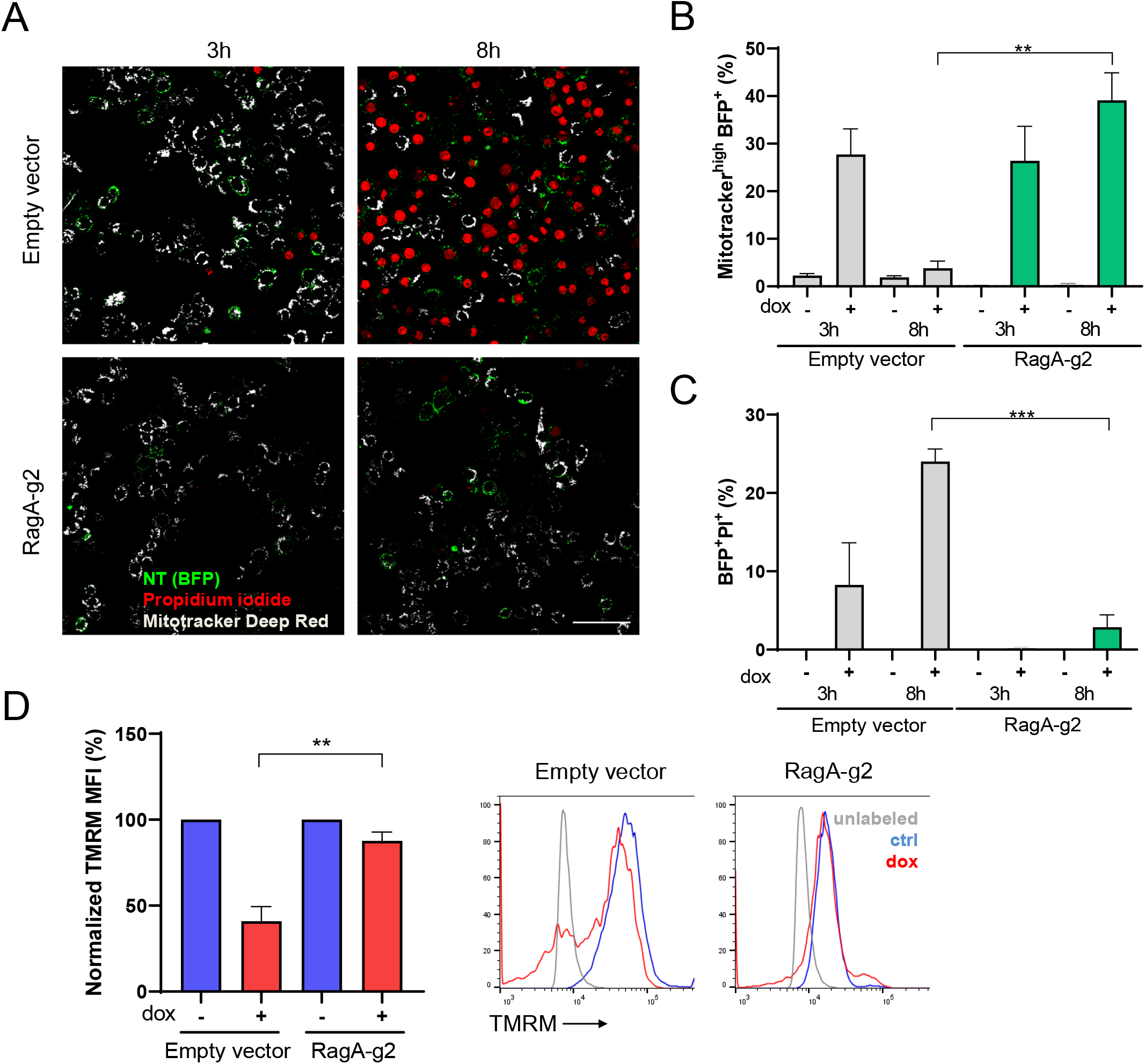
RagA-deficient cells are resistant to pore formation and mitochondrial damage. (A) Representative 246×246 μm^2^ images utilized to analyze effects of NT-GSDMD expression (green) on mitochondria (labeled with Mitotracker Deep Red FM (gray)) and pore formation (PI, red). Bar represents 50 μm. (B,C) After thresholding such images were used to determine the numbers of Mitotracker Deep Red highly fluorescent and BFP positive cells (B) as well as BFP- and PI-positive cells (C). 3-5 frames were analyzed per experimental point. Average values of individual experiments are shown. Mean and SEM of 3 independent experiments are shown. Unpaired two-tailed t-test was used for statistical evaluation. (D) Expression of NT-GSDMD in Empty Vector transduced cells induces a drop in mitochondrial potential as detected by TMRM stain followed by flow cytometry. Representative histograms are shown. Normalized MFI with mean and SEM from 3 individual experiments are shown. MFI of samples 8h after doxycycline addition was normalized to non-treated (100%) and unlabeled (0%) control. Unpaired two-tailed t-test was used for statistical evaluation.

**Supplemental Figure 2:**
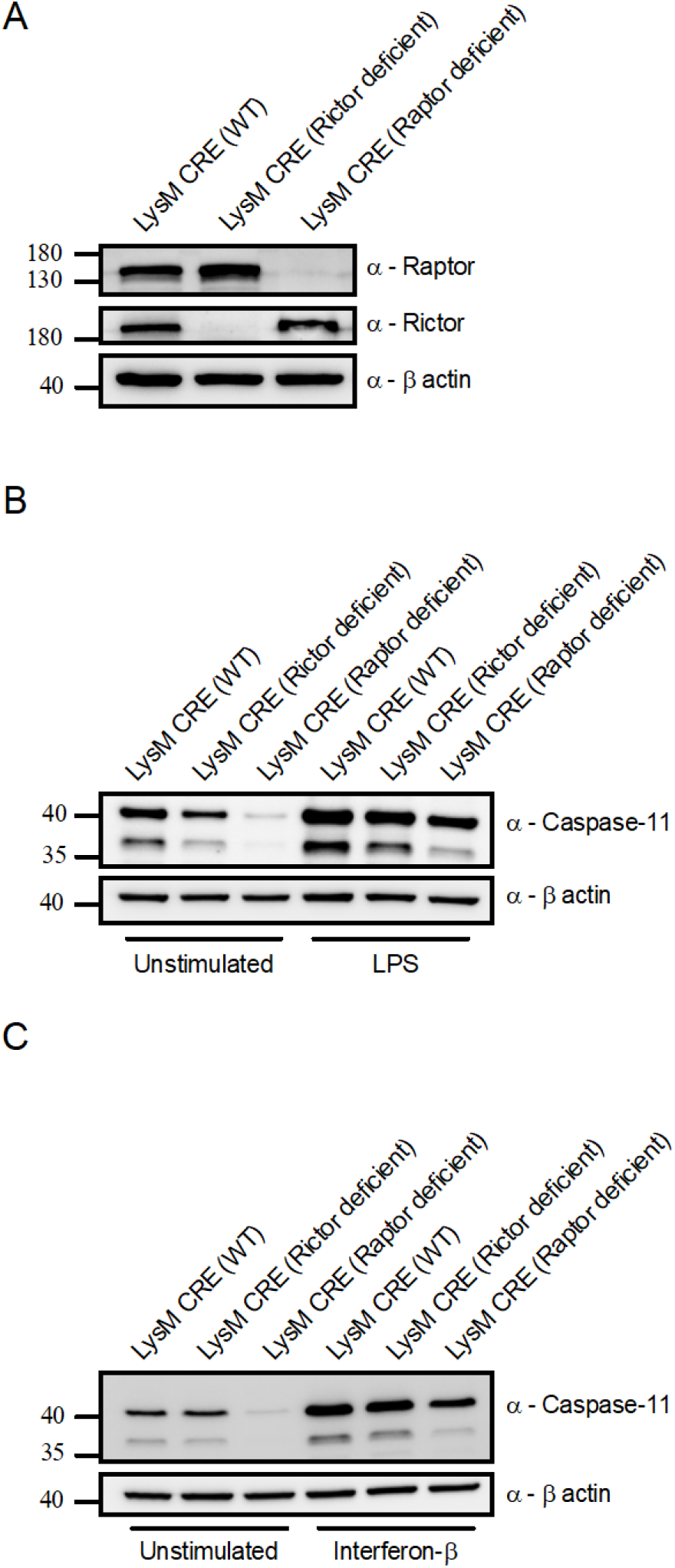
Protein analysis of LysM CRE, Rictor-deficient, and Raptordeficient iBMDMs. (A) Western blot analysis of lysates from unstimulated iBMDMs from LysM CRE, Rictor deficient, and Raptor deficient genotypes investigating protein expression of Raptor, Rictor and β-actin loading control. (B) Western blot analysis of lysates from unstimulated or LPS (1 μg/ml for 3 hours) stimulated iBMDMs from LysM CRE, Rictor deficient, and Raptor deficient genotypes investigating protein expression of caspase-11 and β-actin loading control. (C) Western blot analysis of lysates from unstimulated or IFNβ (1000U for 3 hours) stimulated iBMDMs from LysM CRE, Rictor deficient, and Raptor deficient genotypes investigating protein expression of caspase-11 and β-actin loading control.

## Supplemental Video 1

Related to Figure 3.

Time-lapse live cell imaging of empty vector transduced iBMDMs during 8 hours of doxycycline (2 μg/ml) induction of NT-GSDMD. Left panel is brightfield. Middle panel is propidium iodide fluorescence. Right panel is a merge of these two channels. Images were captured every 30 minutes at 37°C in 5% CO_2_.

## Supplemental Video 2

Related to Figure 3.

Time-lapse live cell imaging of RagA guide 1 transduced iBMDMs (clone C) during 8 hours of doxycycline (2 μg/ml) induction of NT-GSDMD. Left panel is brightfield. Middle panel is propidium iodide fluorescence. Right panel is a merge of these two channels. Images were captured every 30 minutes at 37°C in 5% CO_2_.

## Supplemental Video 3

Related to Figure 3.

Time-lapse live cell imaging of RagA guide 2 transduced iBMDMs (clone C) during 8 hours of doxycycline (2 μg/ml) induction of NT-GSDMD. Left panel is brightfield. Middle panel is propidium iodide fluorescence. Right panel is a merge of these two channels. Images were captured every 30 minutes at 37°C in 5% CO_2_.

## Supplemental Video 4

Related to Figure 5.

Time-lapse live cell imaging of LysM CRE WT iBMDMs primed with IFNβ for 3 hours and electroporated with LPS (1 μg/million cells) for 1 hour following equilibration with propidium iodide containing media. Left panel is brightfield. Middle panel is propidium iodide fluorescence. Right panel is a merge of these two channels. Images were captured every 10 minutes at 37°C in 5% CO_2_.

## Supplemental Video 5

Related to Figure 5.

Time-lapse live cell imaging of LysM CRE Rictor deficient iBMDMs primed with IFNβ for 3 hours and electroporated with LPS (1 μg/million cells) for 1 hour following equilibration with propidium iodide containing media. Left panel is brightfield. Middle panel is propidium iodide fluorescence.

Right panel is a merge of these two channels. Images were captured every 10 minutes at 37°C in 5% CO_2_.

## Supplemental Video 6

Related to Figure 5.

Time-lapse live cell imaging of LysM CRE Raptor deficient iBMDMs primed with IFNβ for 3 hours and electroporated with LPS (1 μg/million cells) for 1 hour following equilibration with propidium iodide containing media. Left panel is brightfield. Middle panel is propidium iodide fluorescence. Right panel is a merge of these two channels. Images were captured every 10 minutes at 37°C in 5% CO_2_.

